# The human inactive X chromosome modulates expression of the active X chromosome

**DOI:** 10.1101/2021.08.09.455676

**Authors:** Adrianna K. San Roman, Alexander K. Godfrey, Helen Skaletsky, Daniel W. Bellott, Abigail F. Groff, Hannah L. Harris, Laura V. Blanton, Jennifer F. Hughes, Laura Brown, Sidaly Phou, Ashley Buscetta, Paul Kruszka, Nicole Banks, Amalia Dutra, Evgenia Pak, Patricia C. Lasutschinkow, Colleen Keen, Shanlee M. Davis, Nicole R. Tartaglia, Carole Samango-Sprouse, Maximilian Muenke, David C. Page

## Abstract

Researchers have presumed that the “inactive” X chromosome (Xi) has little impact, in trans, on the “active” X (Xa). To test this, we quantified Xi and Xa gene expression in individuals with one Xa and zero to three Xi’s. Our linear modeling revealed modular Xi and Xa transcriptomes and significant Xi-driven expression changes for 38% (162/423) of expressed X-chromosome genes. By integrating allele-specific analyses, we found that modulation of Xa transcript levels by Xi contributes to many of these Xi-driven changes (≥ 121 genes). By incorporating metrics of evolutionary constraint, we identified 10 X-chromosome genes most likely to drive sex differences in common disease, and sex chromosome aneuploidy syndromes. We conclude that human X chromosomes are regulated both in cis, through Xi-wide transcriptional attenuation, and in trans, through positive or negative modulation of individual Xa genes by Xi. The sum of cis and trans effects differs widely among genes.

## Introduction

The X chromosome of eutherian mammals exists in two distinct epigenetic states that are referred to as “active” (Xa) and “inactive” (Xi) (Lyon, 1961; 1962; Ohno and Hauschka, 1960). The “n-1” rule (where n is the number of X chromosomes per cell) states that all diploid human somatic cells possess one X chromosome in the active state (Xa), while all other (*i.e.,* n-1) copies of Chr X (Harnden, 1961) are transcriptionally repressed through a mechanism known as X chromosome inactivation (XCI). Despite the name, multiple lines of evidence indicate that Xi is functionally active, making critical contributions to human fitness and viability. For example, 99% of fetuses with only one sex chromosome (45,X) abort spontaneously, suggesting that viability hinges on gene expression from a second sex chromosome – either Xi or Y (Hook and Warburton, 1983; Hook and Warburton, 2014). The rare survivors likely have a mixture of 45,X cells and cells with a second sex chromosome, and they display a constellation of anatomic features known as Turner syndrome (Ford et al., 1959; Turner, 1938).

Studies across four decades have revealed that as many as a quarter of X-linked genes are expressed from Xi in humans; such genes are said to “escape” X inactivation (Balaton et al., 2015). Early studies assayed expression in human-rodent hybrid cell lines that retained Xi but not Xa; detection of Chr X expression in these cells indicates that a gene is expressed from Xi [for example, (Brown et al., 1997; Carrel et al., 1999; Mohandas et al., 1980)]. Subsequent allele-specific methods distinguished transcripts from Xa and Xi in human cell lines with skewed XCI, or in single cells (Carrel and Willard, 2005; Cotton et al., 2013; Garieri et al., 2018; Sauteraud et al., 2021; Tukiainen et al., 2017; Wainer Katsir and Linial, 2019). While conceptually superior to hybrid cell lines, allele-specific methods yielded sparse data because they require the fortuitous presence of heterozygous SNPs to differentiate between alleles. Circumventing this issue, other studies approximated the contributions of Xi to X-linked gene expression by comparing samples with varying Xi copy number, either between 46,XY and 46,XX or between sex chromosome aneuploid and euploid individuals (Craig et al., 2004; Johnston et al., 2008; Nielsen et al., 2020; Raznahan et al., 2018; Sudbrak et al., 2001; Talebizadeh et al., 2006; Trolle et al., 2016; Tukiainen *et al*., 2017; Zhang et al., 2020). These studies employed analytic methods that made it difficult to separate the effect of Xi copy number from the potentially confounding effects of correlated factors like Chr Y copy number, hormonal differences, or tissue composition. More importantly – as underscored by the findings reported here – previous studies assumed the independence and additivity of Xi and Xa expression, without directly testing for influences of one upon the other. In particular, these studies assumed that an increase in expression with additional copies of Xi reflects expression from Xi, which may not always be the case. Given these limitations, we believed that a deeper understanding of Xi gene expression and its consequences was possible.

Here, we used a series of quantitative approaches to investigate gene expression from Xi and Xa. Inspired by previous studies, we took advantage of the natural occurrence in human populations of diverse sex chromosome aneuploidies and performed RNA sequencing (RNA-seq) on two cell types (lymphoblastoid cell lines and primary skin fibroblasts) from 176 individuals spanning 11 different sex chromosome constitutions – from 45,X (Turner Syndrome) to 49,XXXXY. Unlike previous studies, we analyzed the resulting data from all sex chromosome constitutions together – using linear regression models to identify significant changes in Chr X gene expression in identically cultured cells with zero, one, two, or three copies of Xi. Thirty-eight percent of Chr X genes displayed significant Xi-driven expression changes, which we quantified on a gene-by-gene basis using a metric we call ΔE_X_. By combining ΔE_X_ findings with i) allele-specific analyses performed in the same cell lines and ii) published, independent annotations of genes subject to XCI, we discovered that Xi positively or negatively modulates steady-state levels of transcripts of at least 121 genes on Xa, in trans. Thus, Xi and Xa expression are highly interdependent, contrary to the presumptions underlying prior studies. By combining ΔE_X_ with published gene-wise metrics of evolutionary constraint, we identified a set of 10 Chr X genes most likely to drive phenotypes that arise from natural variation in Xi copy number. These 10 candidate “drivers” can now be prioritized in studies of sex differences in common disease, and in explorations of sex chromosome aneuploidy syndromes.

## Results

### Sampling gene expression across sex chromosome constitutions

To conduct a robust, quantitative analysis of Xi’s impacts on X-linked gene expression, we recruited individuals with a wide range of sex chromosome constitutions to provide blood samples and/or skin biopsies (**Fig. 1A**). We generated or received Epstein Barr Virus-transformed B cell lines (lymphoblastoid cell lines, LCLs) and/or primary dermal fibroblast cultures from 176 individuals with one to four X chromosomes and zero to four Y chromosomes. After culturing cells under identical conditions, we profiled gene expression by RNA sequencing (RNA-seq) in LCLs from 106 individuals and fibroblast cultures from 99 individuals (some individuals contributed both blood and skin samples; **Table 1 and Table S1**). To enable analysis at both the gene and transcript isoform levels, we generated 100-base-pair paired-end RNA-seq reads to a median depth of 74 million reads per sample. A resampling (bootstrapping) analysis of our dataset indicates that including more individuals with sex chromosome aneuploidy would only marginally increase the number of differentially expressed genes detected in our analyses (**Fig. S1, Methods**).

**Figure 1.**
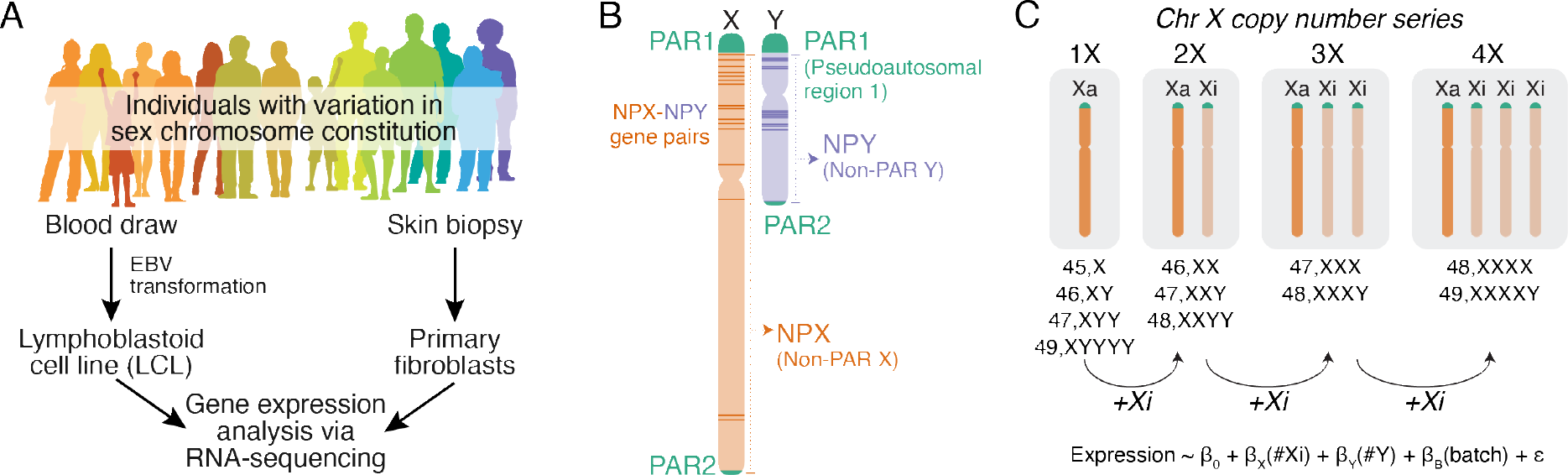
Gene expression analysis of cells from across the spectrum of sex chromosome constitution. **(A)** Collection and processing of samples from individuals with variation in sex chromosome constitution. **(B)** Schematic of the sex chromosomes featuring the X-Y-shared pseudoautosomal regions, PAR1 and PAR2, and the diverged regions, NPX and NPY. **(C)** Linear modeling strategy for analyzing RNA-seq data from individuals with one to four copies of Chr X (zero to three copies of Xi).

**Table 1.** Samples included in sex chromosome aneuploidy analysis.

| Table 1. Samples included in sex chromosome aneuploidy analysis |  |  |
| --- | --- | --- |
| Karyotype | # LCLs | # Fibroblast cultures |
| 45,X | 17 | 23 |
| 46,XX | 22 | 20 |
| 46,XY | 17 | 14 |
| 47,XXX | 7 | 4 |
| 47,XXY | 11 | 30 |
| 47,XYY | 10 | 5 |
| 48,XXXX | 1 | 0 |
| 48,XXXY | 4 | 1 |
| 48,XXYY | 3 | 0 |
| 49,XXXXY | 12 | 1 |
| 49,XYYYY | 2 | 1 |
| Total: | 106 | 99 |

### A metric for the impact of Xi on gene expression

To leverage the full power of our datasets, we collected all of the RNA-seq data for each cell type into a single analysis. We included protein-coding and well-characterized lncRNA genes with a median expression in either 46,XX or 46,XY samples of at least 1 transcript per million (TPM), resulting in 423 Chr X genes analyzed in LCLs and/or fibroblasts. These genes reside within structurally and evolutionarily distinct regions (**Fig. 1B**): two pseudoautosomal regions (PAR1 and PAR2), which are identical in sequence between Chr X and Y, and the non-pseudoautosomal region of the X (NPX), which has diverged in structure and gene content from the non-pseudoautosomal region of the Y [NPY, (Ross et al., 2005; Skaletsky et al., 2003)]. Despite this divergence, 17 homologous “NPX-NPY pair genes” with varying degrees of X-Y similarity in sequence and function remain (Bellott et al., 2014; Skaletsky *et al*., 2003).

We hypothesized that each copy of Xi would incrementally increase expression of some Chr X genes and, therefore, for each gene we modeled expression as a linear function of Xi copy number, controlling for Chr Y copy number and batch (**Fig. 1C**; **Methods**). To assess whether expression of each Chr X gene changed linearly per Xi, we fit non-linear least square regression models to the expression data using power functions (**Methods**). Most NPX and PAR1 genes previously annotated as escaping XCI were best fit by linear models in which expression increases by a fixed amount per Xi, while most genes previously annotated as subject to XCI were best fit by models with no change in expression per Xi (**Fig. S2;** see **Methods** for XCI status annotations from published studies). These results confirm the “n-1” rule at the transcriptomic level, indicating that each cell has a single copy of Xa and n-1 copies of Xi. Moreover, linear modeling revealed that contributions by Xi to Chr X gene expression are strikingly modular, meaning that each Xi is more or less equivalent.

Linear models allowed us to identify genes whose expression changed significantly with additional copies of Xi and to quantify the *absolute* changes in expression (*i.e.*, changes in read counts per Xi). To compare genes expressed at different levels, we also quantified the *relative* changes in expression per Xi. Specifically, we divided the change in expression per Xi (slope of regression, ß_X_) by the expression from the single Xa (average intercept across batches, ß_0_) – a metric we refer to as ΔE_X_ (**Fig. 2A**). ΔE_X_=0 indicates that adding one or more copies of Xi does not affect the level of expression (e.g., *PRPS2*, **Fig. 2B, S3A**); ΔE_X_>0 indicates that expression increases under these circumstances (e.g., *KDM5C*, **Fig. 2C, S3B**), with ΔE_X_=1 suggesting that Xa and Xi contribute equally; and ΔE_X_<0 indicates that expression decreases (e.g., *F8*, **Fig. 2D, S3C**). *XIST,* the long non-coding RNA that acts in *cis* to transcriptionally repress X chromosomes from which it is expressed (Brown et al., 1991; Penny et al., 1996), was the only gene without detectable expression in cells with one copy of Chr X (Xa) that was expressed robustly in cells with a second, inactive copy (Xi). Considering samples with two or more X chromosomes, we found that *XIST* expression increased linearly with each additional copy of Xi (**Fig. 2E, S3B**).

**Figure 2.**
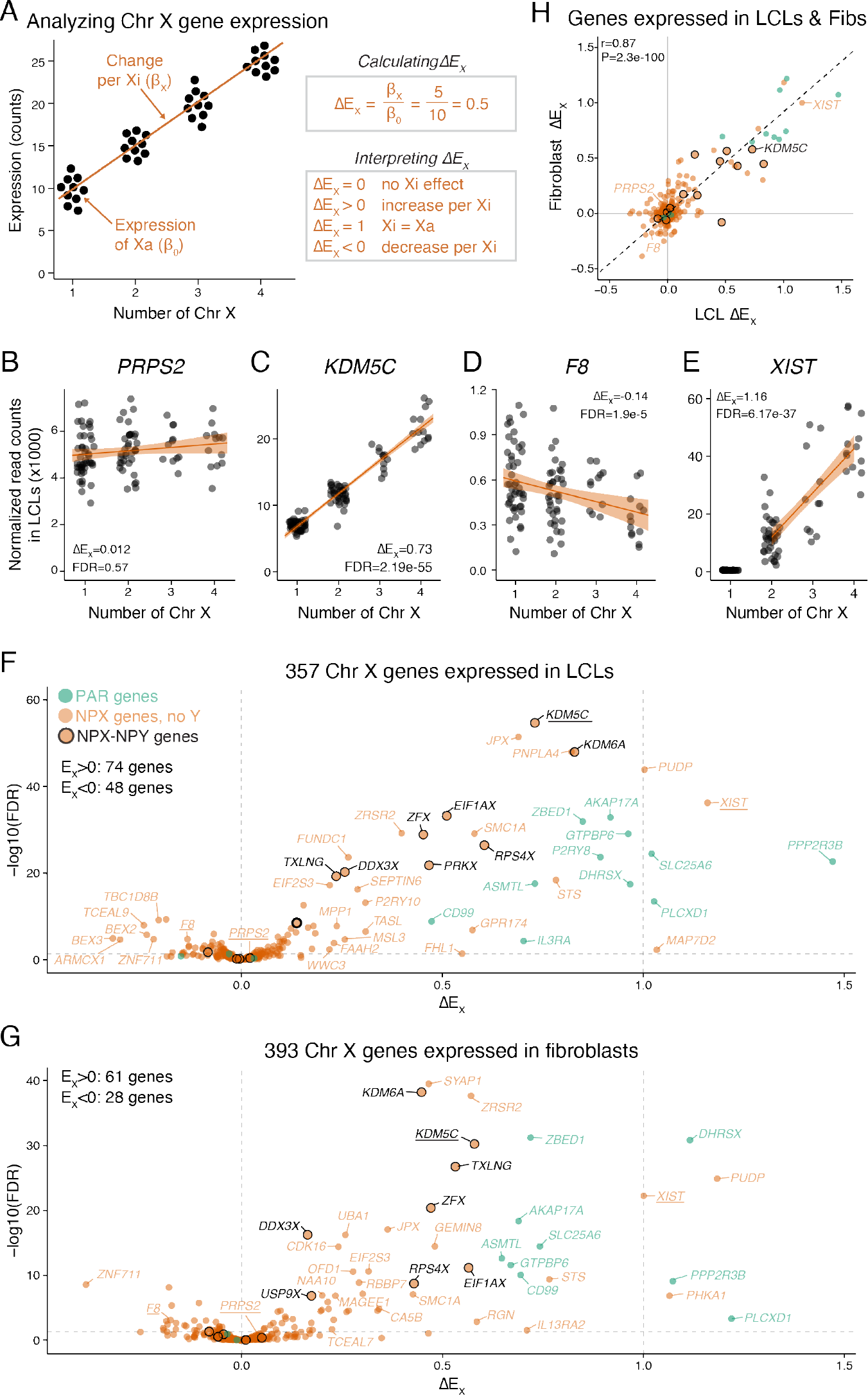
Quantitative assessment of Xi contributions to X chromosome gene expression. **(A)** Schematic scatterplot, linear regression line, and ΔEX calculation for a hypothetical Chr X gene. Each point represents the expression level for an individual sample with the indicated number of copies of Chr X. The calculated coefficients from the linear model in fig. 1C are used to derive ΔEX. **(B-E)** Actual scatterplots and regression lines with confidence intervals for selected Chr X genes in LCLs, representing a range of ΔEX values. Adjusted p-values (FDR) <0.05 indicate that ΔEX values are significantly different from 0. **(F-G)** Scatterplots of ΔEX versus significance for all Chr X genes expressed in LCLs **(F)** and fibroblasts **(G)** illustrate variation in Xi contributions to Chr X gene expression; genes with FDR<0.05 and |ΔEX|≥0.2 are labelled; genes depicted in B-E are underlined. **(H)** Scatterplot comparing ΔEX in LCLs and fibroblasts for 327 Chr X genes expressed in both cell types. Colors as in F-G. Deming regression line and Pearson correlation are indicated.

### ΔE_X_ values vary widely among Chr X genes but not between cell types

Analyzing ΔE_X_ values across Chr X genes revealed that Xi contributions to expression varied widely. Expression of 235 (66%) Chr X genes in LCLs and 304 (77%) in fibroblasts did not change significantly with addition of Xi’s (ΔE_X_≈0), consistent with expression from a cell’s first X chromosome (Xa) and silencing on all others (Xi) (**Fig. 2F-G**; full results in **Table S2**). The remaining 122 (34%) Chr X genes in LCLs and 89 (23%) in fibroblasts had significantly negative or positive ΔE_X_ values. Combining the results in LCLs and fibroblasts, Xi copy number significantly impacts gene expression levels for 38% (162/423) of expressed Chr X genes. NPX genes’ values ranged from −0.39 to 1.2 (**Fig. 2F-G**). PAR1 genes had values near 1, while PAR2 genes had values near zero (**Table S2**). (The stark difference between PAR1 and PAR2 likely reflects their evolutionary origins: PAR1 was preserved on Chr X and Y through sex chromosome evolution and retains autosome-like features, while PAR2 evolved later through a transposition from Chr X to Chr Y (Ciccodicola et al., 2000)). For nearly all Chr X genes, the change in expression per Xi falls short of that contributed by Xa (ΔE_X_<1), similar to previous studies using allelic ratio analysis (Carrel and Willard, 2005; Cotton *et al*., 2013). Only two NPX genes – *XIST* and *PUDP* – and three PAR1 genes – *DHSRX, PLCXD1,* and *PPP2R3B* – showed ΔE_X_ values approaching or exceeding 1 in both LCLs and fibroblasts.

We assessed whether these Chr X expression dynamics were influenced by factors apart from Xi count. We found few differences between cell types; genes expressed in both LCLs and fibroblasts displayed concordant ΔE_X_ values (**Fig. 2H**). This is consistent with studies of differential expression between 46,XY and 46,XX tissues that found correlated expression changes for Chr X genes across diverse tissues (Tukiainen *et al*., 2017). To control for any effects of gonadal sex or Y chromosome copy number on our results, we re-analyzed the data from samples with zero (45,X; 46,XX; 47,XXX; 48,XXXX) or one copy of Chr Y (46,XY; 47,XXY; 48,XXXY; 49,XXXXY), modeling expression as a function of Xi copy number and batch. ΔE_X_ values were unaffected by the presence or absence of a Y chromosome (**Fig. S4**). Because of its design, our study reveals that these consistent Chr X expression dynamics derive from direct, cell-autonomous contributions of Xi rather than systemic effects of hormones or environmental factors.

For genes with multiple transcript isoforms (alternative transcripts), we asked whether ΔE_X_ values were consistent between isoforms (**Methods**). For most genes, transcript isoforms displayed concordant ΔE_X_ values. However, for 33 (19% of) genes with multiple transcript isoforms in LCLs and 25 (13%) in fibroblasts, ΔE_X_ values were discordant: at least one isoform’s ΔE_X_ differed significantly from zero (FDR<0.05) while another isoform’s ΔE_X_ did not (**Fig. S5, Table S3**). The most striking case is that of *UBA1*, where alternative transcription start sites, separated by a CTCF binding site, display divergent behaviors (**note S1**).

To assess reproducibility, we compared our results to those from an independent dataset that used microarrays to assay gene expression in LCLs across diverse sex chromosome constitutions (Raznahan *et al*., 2018). Reanalyzing this dataset using linear models, we found that the resulting microarray ΔE_X_ values correlated well with the ΔE_X_ values calculated from our RNA-seq data (**Fig. S6; Methods**).

### Supernumerary copies of chromosomes Y and 21 show little attenuation of gene expression

To determine whether the attenuated expression observed with extra copies of Chr X also occurs with additional copies of other chromosomes, we analyzed cells from individuals with additional copies of Chr Y or with trisomy 21, a common autosomal aneuploidy and the cause of Down syndrome (Megarbane et al., 2009).

For Chr Y we used the same linear model as for Chr X: modeling expression as a function of Chr Y copy number, Xi copy number, and batch (**Fig. 3A, Methods**). We calculated ΔE_Y_ values separately for NPY and PAR genes, because NPY genes are not expressed in samples with zero Y chromosomes, while PAR genes are expressed from Chr X in all samples.

**Figure 3.**
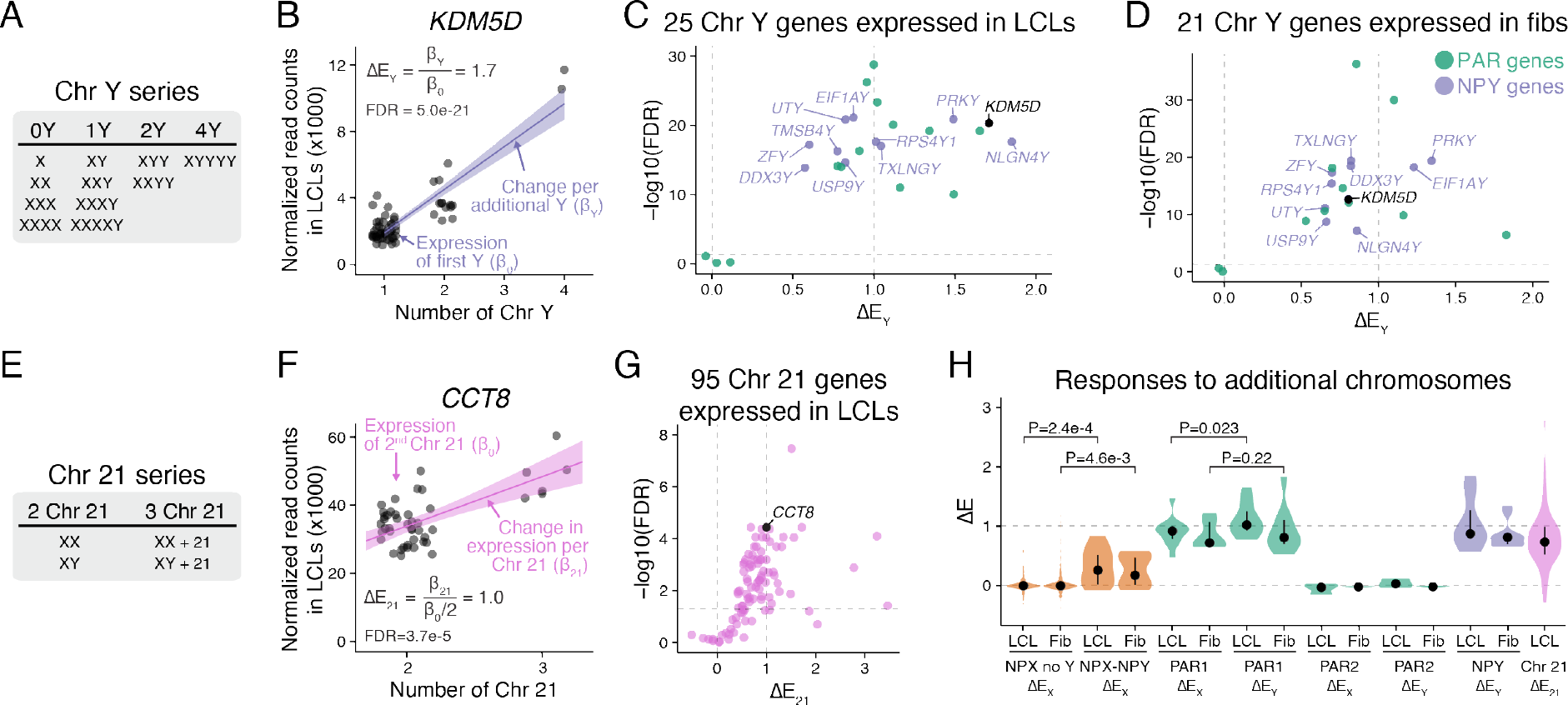
Contributions of Chr Y or 21 copy number to gene expression. **(A)** Chr Y copy number series with zero to four copies. **(B)** Each point shows the expression of NPY gene *KDM5D* in one LCL sample across the Chr Y copy number series, with the regression line and its confidence interval plotted. The formula for calculating ΔEY from the regression coefficients is indicated. **(C-D)** Scatterplot of ΔEY versus significance for all Chr Y genes expressed in LCLs **(C)** or fibroblasts **(D)**; all NPY genes are labelled; *KDM5D,* depicted in B, is shown in black. **(E)** Chr 21 copy number series with two to three copies. **(F)** Each point shows the expression of *CCT8* in one LCL sample across the Chr 21 copy number series, with the regression line and its confidence interval plotted. The formula for calculating ΔE21 from the regression coefficients is indicated. **(G)** Scatterplot of ΔE21 versus significance for all Chr 21 genes expressed in LCLs. *CCT8*, depicted in F, is shown in black. **(H)** Violin plots with median and interquartile range for ΔE values of NPX (without or with a NPY homolog), PAR, NPY, and Chr 21 genes. P-values are listed for comparisons referenced in the text. ΔEX values for NPX genes with and without a Y homolog were compared using Wilcoxon rank sum test. ΔEX and ΔEY values for PAR1 genes wee compared using paired t-test.

For NPY genes, we analyzed samples with one to four Y chromosomes to quantify expression differences, if any, between the first and additional Y chromosomes. Expression of all NPY genes increased significantly, with ΔE_Y_ values close to 1, consistent with near-equal expression from each copy of Chr Y (e.g., *KDM5D*; **Fig. 3B-D, S3D,** full results in **Table S4**).

For PAR genes, we analyzed samples with zero to four Y chromosomes. As with Chr X, PAR1 gene expression increased with additional copies of Chr Y, yielding ΔE_Y_ values close to 1, whereas PAR2 genes had ΔE_Y_ values near zero (**Fig. 3C,D**). This implies, first, that PAR1 genes are expressed on each copy of Chr X or Y, while PAR2 genes are only expressed on the first copy of Chr X (Xa); and second, that PAR1 gene expression from each additional Chr X or Y is roughly equal to expression from the first.

Finally, we examined Chr 21 gene expression as a function of Chr 21 copy number (**Fig. 3E**; **Methods**). Nearly three-quarters of expressed Chr 21 genes significantly (FDR<0.05) increased in expression with an additional copy of Chr 21 (e.g., *CCT8*; **Fig. 3F**), and none significantly decreased (**Fig. 3G; Table S5**). These results align well with independent studies of Chr 21 gene expression (Sullivan et al., 2016).

In sum, our analysis reveals that most genes on Chr Y and Chr 21 are expressed similarly on each copy of their respective chromosomes. The median ΔE values for Chr Y (including NPY and PAR1) and Chr 21 genes range from 0.74 to 1.0. By comparison, NPX genes without NPY homologs had median ΔE_X_≈0, while NPX genes with NPY homologs had modestly higher median ΔE_X_ values (LCLs: 0.26, fibroblasts: 0.17; **Fig. 3H**). Even PAR1 genes, which as a group had the highest median ΔE_X_ values, were modestly attenuated on Xi compared to Chr Y, especially in LCLs (**Fig. 3H**). This Y-vs-X effect was most pronounced for *CD99*, located near the PAR1-NPX/Y boundary (**Tables S2 and S4**), consistent with suggestions that PAR1 gene expression on Xi is modestly attenuated by spreading of heterochromatin (Tukiainen *et al*., 2017). These differences highlight the absence of a chromosome-wide mechanism attenuating (or otherwise altering) gene expression on supernumerary copies of Chr Y and Chr 21, in contrast to Chr X.

### Xi modulation of Xa transcript levels revealed by divergence of ΔE_X_ and allelic ratio

ΔE_X_ conveys the change in a gene’s expression due to an additional Xi regardless of the mechanism(s) responsible for this change. We hypothesized that a Chr X gene’s ΔE_X_ value could reflect the combined effects of two mechanisms: 1) transcription of Xi allele(s) and 2) modulation of the Xa allele by Xi in trans.

Seeking evidence of these mechanisms, we searched gene by gene for agreements and disagreements between our calculated ΔE_X_ values and published descriptions of the genes as “escaping” XCI (being expressed from Xi) or being subject to it (silenced on Xi). For this purpose, we curated annotations of each expressed gene’s XCI status from studies of allele-specific expression (Carrel and Willard, 2005; Cotton *et al*., 2013; Garieri *et al*., 2018; Sauteraud *et al*., 2021; Tukiainen *et al*., 2017) (**Methods; Table S6A**). Many genes with ΔE_X_>0 were classified as expressed from Xi (50/74 in LCLs and 43/61 in fibroblasts), indicating that transcription from Xi alleles underlies their ΔE_X_ values (**Fig. 4A; Table S6A**). Genes with these characteristics overlapped significantly between fibroblasts and LCLs (**Fig. 4B**).

**Figure 4.**
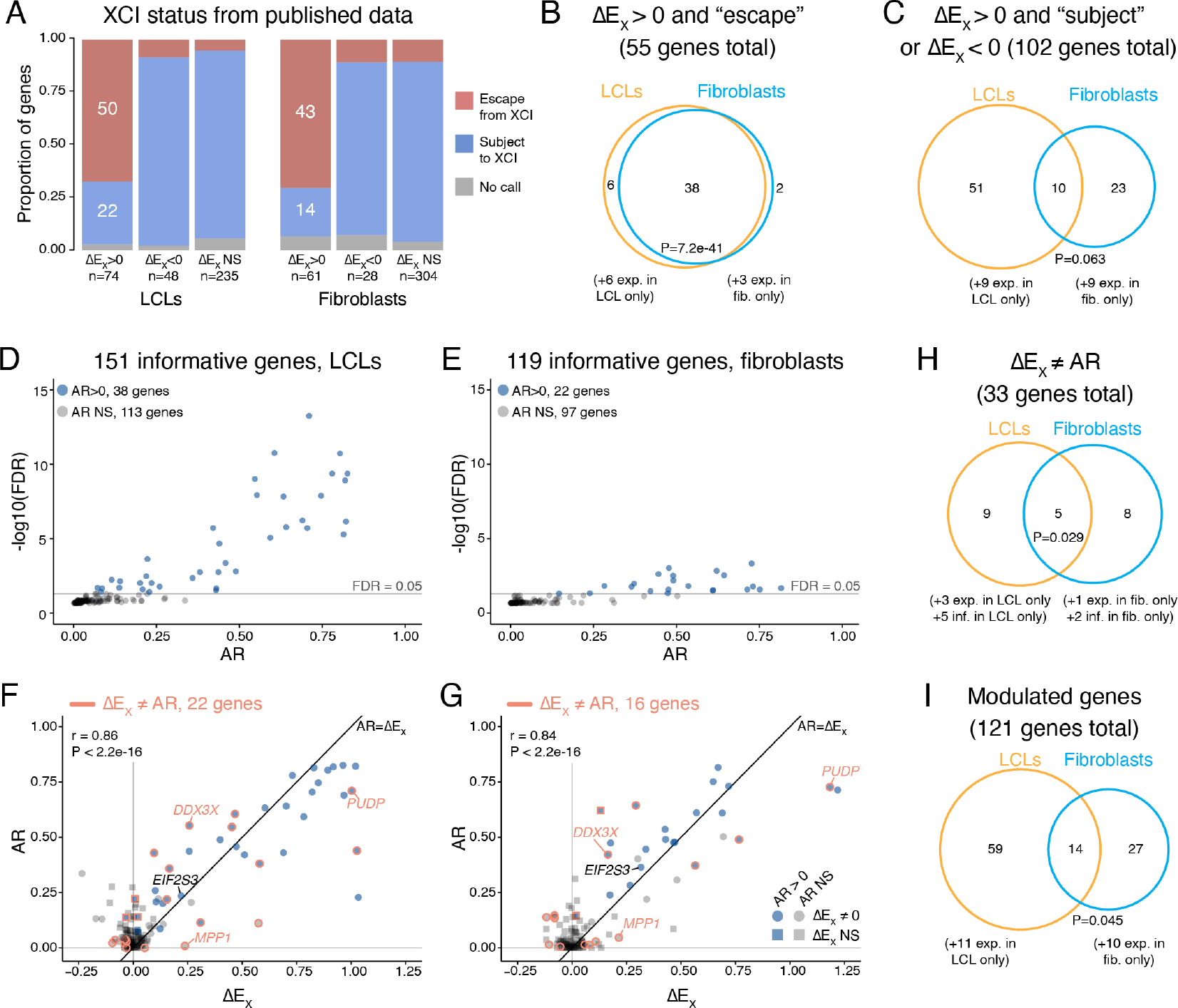
Comparison of ΔEX values with allelic ratios (AR) reveals that Xi modulates Xa expression. **(A)** Stacked barplots for genes with ΔEX values greater than, less than, or approximately equal to zero, apportioned by their annotated XCI status from published studies (see **Methods** and **Table S6A** for newly-compiled XCI status calls). **(B-C)** Venn diagrams comparing LCLs and fibroblasts for genes with ΔEX values that are either **(B)** explained or **(C)** not explained by published XCI status. Genes expressed in both cell types were included in the Venn diagrams, and genes with cell-type specific expression are noted below. **(D-E)** Each point shows the mean adjusted AR for an informative gene (with heterozygous SNPs in at least 3 samples with skewed XCI) and whether AR is significantly greater than zero in **(D)** LCLs or **(E)** fibroblasts. **(F-G)** Each point denotes AR and ΔEX values for an AR-informative gene in **(F)** LCLs or **(G)** fibroblasts. In **(F-G)**, the color of the point indicates whether the gene’s AR value is significantly greater than zero (blue) or not (grey); the shape indicates whether the gene’s ΔEX value is significantly different from zero (circles) or not (squares); and an orange outline indicates that ΔEX differs significantly from AR. Black diagonal line, AR = ΔEX. Pearson correlation coefficients (r) and P-values are indicated. **(H)** Venn diagram comparing LCLs and fibroblasts for genes with ΔEX values not equal to their AR values. Genes expressed and informative in both cell types are depicted in the Venn diagram, with genes that are cell-type specific or informative in only one cell type indicated below. **(I)** Venn diagram comparing all modulated genes in LCLs and fibroblasts (the union of figures **(C)** and **(H)**). All Venn diagram P-values, hypergeometric test.

For 102 (24%) of the 423 Chr X genes we evaluated in LCLs or fibroblasts, our calculated ΔE_X_ values were at odds with expectations arising from the published annotations of XCI status (**Fig. 4A,C; Table S6A**). For example, among genes with ΔE_X_>0, 22 in LCLs and 14 in fibroblasts were described as silenced on Xi. Additionally, previous models offer no explanation for the 48 genes in LCLs and 28 in fibroblasts with ΔE_X_<0, most of which were described as silenced on Xi (**Fig. 4A,C; Table S6A**). Genes with these characteristics did not overlap significantly between LCLs and fibroblasts, even though most are expressed in both cell types, indicating that this regulation is largely cell-type-specific (**Fig. 4C**). These unanticipated findings are unlikely to reflect experimental error in the previous or current studies. Instead, they suggest that, for many Chr X genes whose Xi allele(s) are silent, the Xa allele is nonetheless up-regulated (ΔE_X_>0) or down-regulated (ΔE_X_<0) by Xi.

To corroborate these findings in our own dataset, we performed an allele-specific analysis in our LCL and fibroblast samples with two X chromosomes. To distinguish between Chr X alleles, we identified heterozygous single nucleotide polymorphisms (SNPs) in expressed genes (**Methods**). We then identified samples in our dataset with skewed XCI (21 LCL and 10 fibroblast samples; see **note S2**), and used these samples to compute the average ratio of Xi to Xa expression (the allelic ratio, AR) for each gene. To calculate the AR, we required heterozygous SNPs in at least three samples, resulting in AR values for 151 genes in LCLs and 119 in fibroblasts (**Table S6**). In LCLs and fibroblasts, respectively, 38 (25%) and 22 (18%) of these genes had AR values significantly greater than zero, indicating that they are expressed from Xi (**Fig. 4D-E**); these results agreed well with published AR values (**note S2**).

We next compared each gene’s AR and ΔE_X_ values. If Xi and Xa expression are fully independent of each other, and therefore additive, we would expect the AR for a given gene to approximate its ΔE_X_ value. However, if Xi modulates the gene’s Xa transcript levels in trans, then independence and additivity will not be observed, and instead the gene’s ΔE_X_ and AR values will differ. Most X-linked genes, e.g., *EIF2S3*, had AR values that approximate their ΔE_X_ values, and AR and ΔE_X_ were highly correlated among many informative genes in both LCLs and fibroblasts (**Fig. 4F-G**). For these genes, the ΔE_X_ value may directly reflect the level of transcription from Xi.

However, for 33 informative genes in LCLs or fibroblasts, AR and ΔE_X_ were significantly different, indicating that Xi regulated Xa transcript levels upward or downward in trans (**Fig. 4F-H**). Some genes, like *MPP1,* were not expressed from Xi (AR≈0), but nonetheless had ΔE_X_ values significantly different from zero: 0.24 in LCLs and 0.21 in fibroblasts, indicating that levels of Xa-derived transcripts are positively regulated by Xi. Other genes, like *DDX3X* and *PUDP*, had significant expression from Xi (AR>0, FDR<0.05) *and* evidence of Xi regulation of steady-state expression levels. *DDX3X* had an AR (LCLs: 0.55, fibroblasts: 0.42) that is significantly higher than its ΔE_X_ value (LCLs: 0.26, fibroblasts: 0.16) in both LCLs and fibroblasts, indicating both that *DDX3X* is expressed on Xi *and* that its steady-state transcript levels are negatively regulated by Xi. Conversely, *PUDP* had an AR (LCLs: 0.71, fibroblasts: 0.73) that is significantly lower than its ΔE_X_ value (LCLs: 1.0, fibroblasts: 1.2), indicating both that *PUDP* is expressed on Xi *and* that the gene’s steady-state transcript levels are positively regulated by Xi.

These analyses, combining ΔE_X_ with published or newly derived AR data, provide a rich portrait of X-linked gene regulation. Xi can impact expression levels of an X-linked gene through two mechanisms: transcription of the Xi allele and modulation of steady-state transcript levels by Xi in trans. These mechanisms can operate independently of each other, or together, on a gene-by-gene basis, and each of the two mechanisms affects a sizeable fraction of all X-chromosome genes. Of 423 X-chromosome genes expressed in LCLs and/or fibroblasts, at least 121 genes (29%) are modulated on Xa by Xi in one or both cell types (**Fig. 4I**). (This represents the union of the 102 genes for which the public AR data cannot explain the ΔE_X_ values (**Fig. 4C**) and the 33 genes with AR values significantly different from ΔE_X_ (**Fig. 4H**).) The observed modulation of steady-state transcript levels suggests that Xi regulates the expression of genes on Xa in trans.

### Combining ΔE_X_ and expression constraint metrics identifies likely drivers of Xi-associated phenotypes

While the somatic cells of all diploid individuals have one Xa, the number of Xi’s varies in the human population from zero to four. This variation is associated with many important differences in phenotypes and disease predispositions, for example, those observed between 45,X (Turner syndrome) and 46,XX individuals, between 47,XXY (Klinefelter syndrome) and 46,XY individuals, or even between 46,XY males and 46,XX females. We hypothesized that phenotypes and predispositions associated with Xi copy number are due to changes in the copy numbers of some of the Chr X genes where we found positive or negative ΔE_X_ values. We reasoned that phenotypically critical genes would be “dosage sensitive”, *i.e.*, their expression levels would be tightly constrained by natural selection, while the expression levels of genes whose dosage is not phenotypically critical could vary with little consequence.

To gauge the constraints that selection has imposed on each gene’s expression level, we turned to metrics derived from population and evolutionary genetic studies. We assessed tolerance of under-expression using 1) LOEUF, the ratio of observed to expected loss-of-function (LoF) variants in human populations (Karczewski et al., 2020)), 2) RVIS, the residual variation intolerance score (Petrovski et al., 2013), and 3) pHI, the probability of haploinsufficiency (Huang et al., 2010). Both LOEUF and RVIS use large-scale human genomic sequencing data to evaluate selection against LoF variants, while pHI is based on evolutionary and functional metrics. LoF variants should be culled from the population in genes whose under-expression is deleterious, while they may accumulate in genes whose under-expression has little effect on fitness.

To assess tolerance of over-expression, we examined conservation of targeting by microRNAs (miRNAs; P_CT_ score (Friedman et al., 2009)), which repress expression by binding to a gene’s 3’ untranslated region (Bartel, 2009). Genes sensitive to over-expression have maintained their miRNA binding sites across vertebrate evolution, while genes whose over-expression has little or no effect on fitness show less conservation of these sites (Naqvi et al., 2018).

To weigh these four metrics simultaneously, we calculated each gene’s percentile rank for each metric, from most constrained (high percentile) to least constrained (low percentile). We calculated percentiles separately for autosomal (including PAR1) and NPX genes and then, for each gene, averaged percentile rankings across the four metrics.

We first examined expression constraints for PAR1 genes, whose high ΔE_X_ values suggested they may drive phenotypes associated with Xi copy number. Compared to autosomal genes, PAR1 genes are less constrained on average (P=5.5e-5, Wilcoxon rank sum test), with most ranking in the least-constrained quartile (**Fig. 5A,B, Table S7A**). This indicates that altering their expression levels has little impact on human fitness. Indeed, homozygous LoF mutations have been reported for 3 of 15 PAR1 genes, demonstrating dispensability (Karczewski *et al*., 2020). Only two PAR1 genes, *SHOX* and *SLC25A6,* rank in the more constrained half of the comparison group (**Table 2**). *SHOX* copy number contributes to variation in height in individuals with sex chromosome anomalies (Clement-Jones et al., 2000; Fukami et al., 2016; Ogata and Matsuo, 1993; Ottesen et al., 2010; Rao et al., 1997), while *SLC25A6* has not yet been linked to any phenotype. Apart from these two genes, the high tolerance of under- and over- expression for most PAR1 genes argues against prominent roles in phenotypes associated with Chr X (or X+Y) copy number.

**Figure 5.**
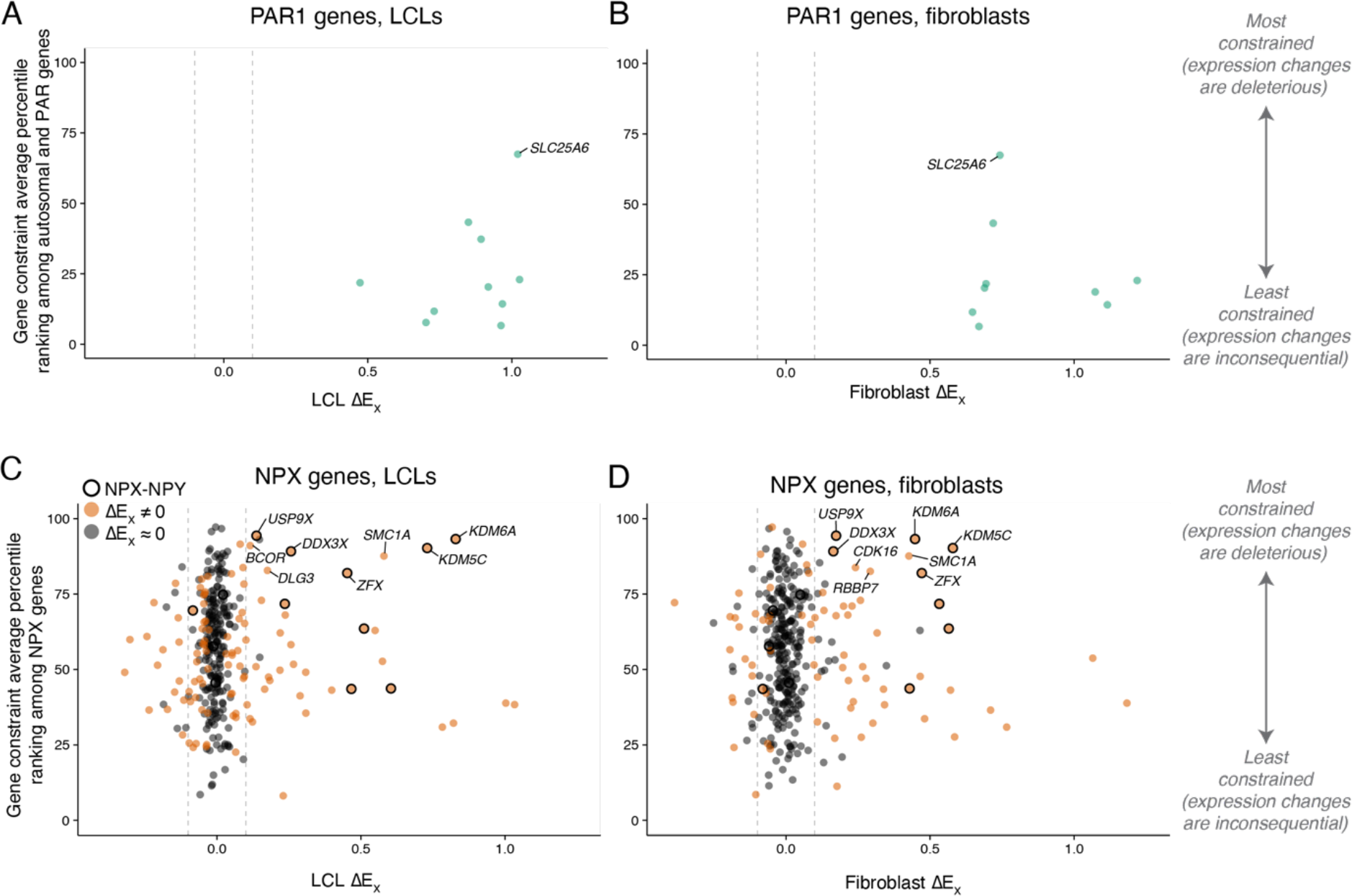
Combining ΔEX with metrics of constraint on expression levels identifies genes likely to contribute to phenotypes associated with Xi copy number. Scatterplots of ΔEX versus gene constraint percentile ranking for PAR1 **(A-B)** or NPX **(C-D)** genes. Each point represents an expressed gene with scores for at least two of the four expression constraint metrics evaluated, excluding ampliconic genes. Dashed lines indicate |ΔEX| thresholds of 0.1 for genes to be considered likely contributors to phenotypes driven by Xi copy number; labeled genes include **(A-B)** *SLC25A6*, the only PAR1 gene to score above the 50^th^ percentile for autosomal and PAR genes and **(C-D)** among NPX genes with |ΔEX| > 0.1, the 10 genes with the highest constraint percentile rankings in LCLs or fibroblasts.

**Table 2.**
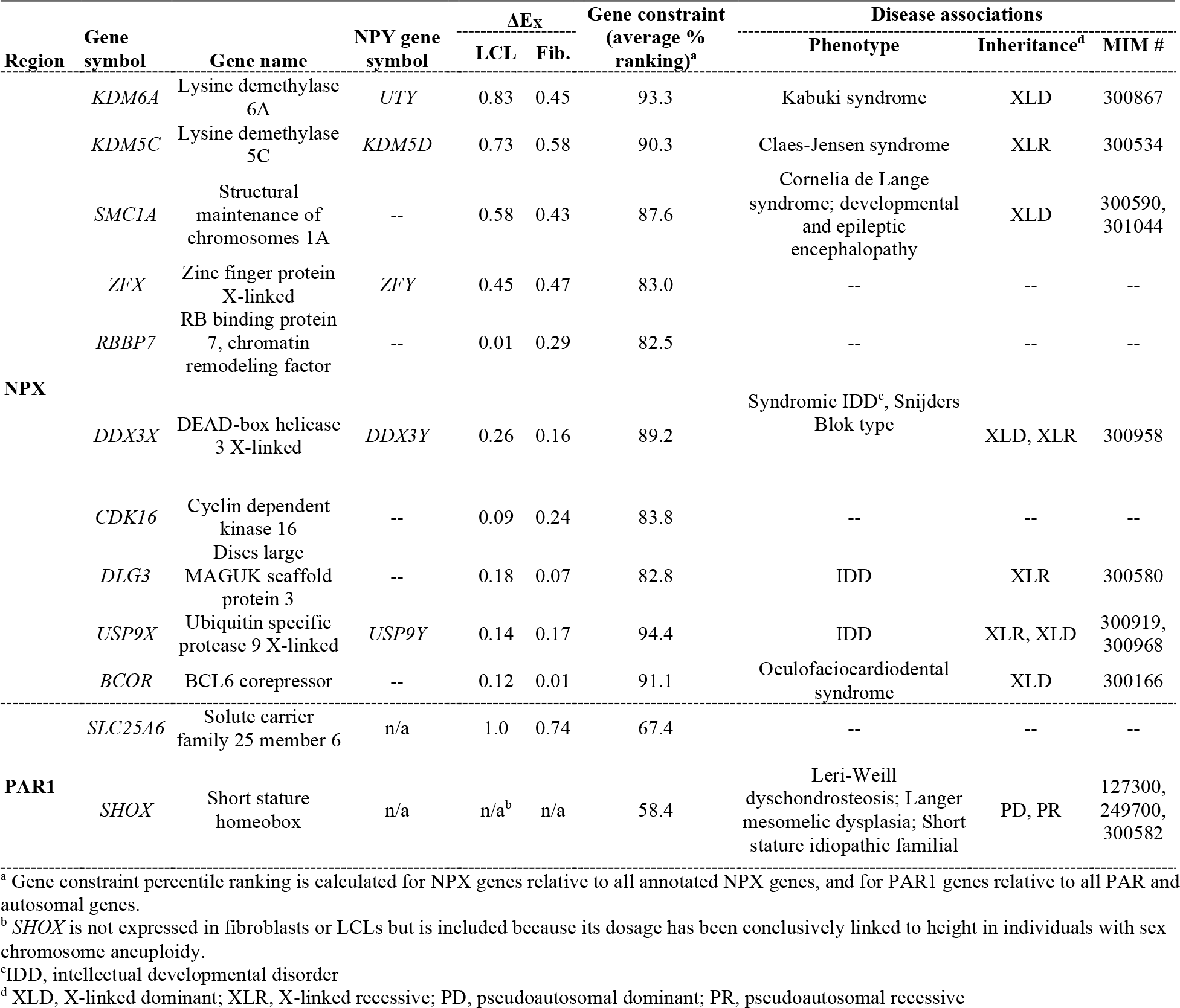
X chromosome genes that may drive the phenotypic impacts of variation in Xi copy number.

| Region | Gene symbol | Gene name | NPY gene symbol | $\Delta E_x$ | | Gene constraint (average % ranking)<sup>a</sup> | Disease associations | | |
| --- | --- | --- | --- | --- | --- | --- | --- | --- | --- |
|  |  |  |  | LCL | Fib. |  | Phenotype | Inheritance<sup>d</sup> | MIM # |
| NPX | *KDM6A* | Lysine demethylase 6A | *UTY* | 0.83 | 0.45 | 93.3 | Kabuki syndrome | XLD | 300867 |
|  | *KDM5C* | Lysine demethylase 5C | *KDM5D* | 0.73 | 0.58 | 90.3 | Claes-Jensen syndrome | XLR | 300534 |
|  | *SMC1A* | Structural maintenance of chromosomes 1A | -- | 0.58 | 0.43 | 87.6 | Cornelia de Lange syndrome; developmental and epileptic encephalopathy | XLD | 300590, 301044 |
|  | *ZFX* | Zinc finger protein X-linked | *ZFY* | 0.45 | 0.47 | 83.0 | -- | -- | -- |
|  | *RBBP7* | RB binding protein 7, chromatin remodeling factor | -- | 0.01 | 0.29 | 82.5 | -- | -- | -- |
|  | *DDX3X* | DEAD-box helicase 3 X-linked | *DDX3Y* | 0.26 | 0.16 | 89.2 | Syndromic IDD<sup>c</sup>, Snijders Blok type | XLD, XLR | 300958 |
|  | *CDK16* | Cyclin dependent kinase 16 | -- | 0.09 | 0.24 | 83.8 | -- | -- | -- |
|  | *DLG3* | Discs large MAGUK scaffold protein 3 | -- | 0.18 | 0.07 | 82.8 | IDD | XLR | 300580 |
|  | *USP9X* | Ubiquitin specific protease 9 X-linked | *USP9Y* | 0.14 | 0.17 | 94.4 | IDD | XLR, XLD | 300919, 300968 |
|  | *BCOR* | BCL6 corepressor | -- | 0.12 | 0.01 | 91.1 | Oculofaciocardiodental syndrome | XLD | 300166 |
| PAR1 | *SLC25A6* | Solute carrier family 25 member 6 | n/a | 1.0 | 0.74 | 67.4 | -- | -- | -- |
|  | *SHOX* | Short stature homeobox | n/a | n/a<sup>b</sup> | n/a | 58.4 | Leri-Weill dyschondrosteosis; Langer mesomelic dysplasia; Short stature idiopathic familial | PD, PR | 127300, 249700, 300582 |

<sup>a</sup> Gene constraint percentile ranking is calculated for NPX genes relative to all annotated NPX genes, and for PAR1 genes relative to all PAR and autosomal genes.
<sup>b</sup> *SHOX* is not expressed in fibroblasts or LCLs but is included because its dosage has been conclusively linked to height in individuals with sex chromosome aneuploidy.
<sup>c</sup> IDD, intellectual developmental disorder
<sup>d</sup> XLD, X-linked dominant; XLR, X-linked recessive; PD, pseudoautosomal dominant; PR, pseudoautosomal recessive

Turning to the much larger set of NPX genes, we found that their widely ranging constraint metrics correlated poorly with their ΔE_X_ values (**Fig. 5C,D, Table S7B**). Thus, ΔE_X_ alone does not predict dosage sensitivity among NPX genes. To identify the NPX genes most likely to drive Xi-copy-number-dependent phenotypes, we selected those with |ΔE_X_| ≥ 0.1 (FDR<0.05) in LCLs or fibroblasts, and ranked these by their average constraint metrics. Five of the top 10 genes by these criteria (**Table 2**) had NPY homologs, a significant enrichment (P=7.0e-4, hypergeometric test), and all five had ΔE_X_ values > 0.1 in both cell types. Of the five genes without NPY homologs, only two had ΔE_X_ values significantly greater than zero in both cell types: *SMC1A* and *CDK16.* The remaining three genes had ΔE_X_ values significantly different from zero in only one of the two cell types analyzed, and they tended to have lower absolute ΔE_X_ values.

If these 10 genes are dosage-sensitive drivers of Xi-dependent phenotypes – even when harboring no mutations – then one might expect mutant phenotypes to be pronounced and to display distinctive modes of inheritance. Accordingly, we searched OMIM for disease annotations. Germline mutations in seven of the 10 genes are reported to cause severe developmental disorders, including well characterized childhood syndromes for five of the genes (**Tables 2, S7B**). Indeed, five of the seven mutation-bearing genes are reported to display dominant inheritance (affected heterozygous females) – a significant enrichment among X-linked genes (P-value = 0.0037, hypergeometric test) and consistent with extraordinary dosage sensitivity. Extrapolating from these findings, we speculate that some of the genes without annotated OMIM phenotypes may have important roles in disease; in the case of *ZFX*, no LoF mutations are reported in gnomAD even though the gene’s roles in regulating stem cell self-renewal and cancer cell proliferation are well documented (Fang et al., 2014; Galan-Caridad et al., 2007; Zhou et al., 2011). Taken together, these 10 genes represent good candidates for driving Xi-dependent phenotypes characteristic of individuals with sex chromosome aneuploidies – as well as differences in disease risks between ordinary (euploid) females and males.

## Discussion

We analyzed Chr X gene expression quantitatively, in two types of cells cultured from individuals with one to four X chromosomes, e.g., 45,X to 49,XXXXY (**Fig. 1**). Folding this diversity of sex chromosome constitutions into a single linear model (**Fig. 2**) yielded advantages over previous studies, which compared sex chromosome constitutions in pairwise fashion, most frequently 45,X vs 46,XX; or 46,XY vs 47,XXY; or 46,XX vs 46,XY. First, our linear model embodied, tested, and confirmed – at the level of the X transcriptome – the so-called “n-1” rule (Harnden, 1961) whereby diploid somatic cells with a given number (n=1, 2, 3, 4) of X chromosomes have a single Xa and n-1 Xi’s. Second, linear modeling provided the power needed to detect and precisely quantify increases or decreases in expression of individual Chr X genes as a function of Chr X copy number. Third, linear modeling revealed that the expression contributions made by each copy of Xi are modular, indicating that each copy of Xi is equivalent, or nearly so, even among unrelated individuals. Fourth, by comparing samples that vary in Xi copy number with and without a Y chromosome, we found that expression from Xa is quantitatively indistinguishable in phenotypic males and females – as is expression from Xi (**Fig. S4**). Thus, both Xi and Xa make modular contributions to Chr X gene expression – contributions independent of and unaffected by the presence of the NPY or the gonadal sex of the individual.

Finally, linear modeling of gene expression as a function of Chr X copy number yielded the metric ΔE_X_, which captures the positive or negative impact of Xi(s) on steady state transcript levels for each gene, normalized to account for gene-to-gene variation in expression level (**Fig. 2A**). Fully 38% (162/423) of expressed Chr X genes in LCLs or fibroblasts displayed a statistically significant positive or negative ΔE_X_ value, indicating that their expression is impacted by the presence of one or more Xi’s. This is nearly double what would be expected based on the prior literature’s estimates of escape, which – even under the broadest definition of escape – includes only 20% (86/423) of expressed Chr X genes (**Table S6A**). ΔE_X_ values varied widely among Chr X genes, from −0.39 to +1.2 (**Fig. 2F-G**), but showed much less variation between the two cell types studied (**Fig. 2H**), suggesting the possibility that the ΔE_X_ “settings” for each gene were established prior to the embryonic divergence of the hematopoietic and skin fibroblast lineages, and subsequently maintained through development.

We extended the utility of the ΔE_X_ metric by cross-referencing and comparing it, one gene at a time, with an orthogonal metric: the allelic ratio (AR) of Xi and Xa transcripts in cells with skewed XCI and SNP heterozygosity. AR values significantly greater than zero unambiguously identify Chr X genes that are expressed from both Xi and Xa (and therefore “escape” XCI) (Carrel and Willard, 1999). By comparing ΔE_X_ and AR values, we discovered that Xi up- or down-modulates Xa expression of at least 121 genes, or nearly 29% of the 423 Chr X genes that are demonstrably expressed in either LCLs or fibroblasts. This modulation is manifest whenever a gene’s AR and ΔE_X_ values differ significantly, and it is most starkly apparent when the gene is not expressed from Xi (*i.e.*, when AR approximates zero) but nonetheless displays a significantly positive or negative ΔE_X_ value. While “escape” from XCI has been well documented over the past four decades (Carrel *et al*., 1999; Carrel and Willard, 2005; Cotton *et al*., 2013; Tukiainen *et al*., 2017), the novel combination of the AR and ΔE_X_ metrics reported here was required to observe modulation, explaining why it has previously been unappreciated.

Thus, combined analysis of ΔE_X_ and AR reveals a nuanced, gene-by-gene tapestry of Xi-driven changes in expression of Chr X genes. For some genes, ΔE_X_ was explained entirely by expression from the Xi allele, while for others ΔE_X_ was explained entirely by modulation – positive or negative – of steady-state RNA levels derived from the Xa allele. For a third set of genes, ΔE_X_ was explained by the combined effects of expression from Xi *and* modulation of steady-state RNA levels. Proposals of uniform, chromosome-wide “X chromosome upregulation” (XCU) during mammalian development or evolution (Ohno, 1967; Okamoto et al., 2021) will need to be revisited in light of this unforeseen diversity of gene-by-gene responses to variation in Chr X copy number.

Finally, we paired the ΔE_X_ metric with population and evolutionary measures of constraint on expression levels to identify 10 NPX genes that are most likely (among the 423 Chr X genes expressed in LCLs and/or fibroblasts) to drive Xi-associated phenotypes (**Fig. 5C,D** and **Table 2**). Despite their high ΔE_X_ values, most PAR (pseudoautosomal) genes did not exhibit the constraints on expression levels that we required for inclusion in this select group of candidate drivers (**Fig. 5A,B**). We propose the 10 NPX genes – five of which have divergent NPY homologs – as potential drivers of i) differences in health and disease between 46,XY and 46,XX cohorts and ii) the distinctive phenotypes associated with sex chromosome aneuploidies, including Turner syndrome (45,X) and Klinefelter syndrome (47,XXY). We speculate that one or more of these 10 NPX genes, which include transcriptional and epigenetic regulators, may also drive the modulation of Xa genes by Xi.

### Limitations of this study

The human individuals sampled here are largely of European ancestry; it will be important to validate these findings in a more ancestrally diverse set of individuals. Our findings in LCLs and fibroblasts were largely concordant, but they may not generalize to all somatic tissues and cell types. Our study focused on 423 Chr X genes that are expressed in LCLs and/or fibroblasts; our conclusions may not generalize to Chr X genes that are not expressed in these cell types. Our list of Chr X genes likely to drive Xi-dependent phenotypes is incomplete, as it is biased toward genes expressed in LCLs and fibroblasts, and toward genes with long ORFs well suited to expression constraint analysis; future studies will add to this list.

## Acknowledgements

We thank members of the Page laboratory, especially Lukas Chmatal, for helpful comments on the manuscript, Sahin Naqvi for advice on RNA-sequencing and analysis, and Jorge Adarme and Susan Tocio for laboratory support. We thank Clare Malone for advice in optimizing the CRISPRi system and the Whitehead Institute Genome Technology Core facility for library preparation and sequencing.

## Funding

National Institutes of Health grant F32HD091966 (AKSR)

National Institutes of Health grant U01HG0007587 (DCP and MM)

National Institutes of Health grant K23HD092588 (SMD)

Schmidt Science Fellows (AFG)

Lallage Feazel Wall Damon Runyon Cancer Research Foundation Fellowship (LVB)

Howard Hughes Medical Institute (DCP)

National Human Genome Research Institute Intramural Research Program (MM)

NIH/NCATS Colorado CTSA grant UL1 TR002535 (NRT)

Contents are the authors’ sole responsibility and do not necessarily represent official NIH views.

Philanthropic gifts from:

Brit and Alexander d’Arbeloff

Arthur W. and Carol Tobin

Brill Matthew Brill

Charles Ellis

## Author contributions

Conceptualization: AKSR and DCP

Data curation: AKSR and HS

Formal analysis: AKSR, AKG, HS, DWB, AG, HLH, LVB, and JFH

Funding acquisition: AKSR, AFG, LVB, SMD, NRT, MM, and DCP

Investigation: AKSR, SP, LB, AD, and EP

Methodology: AKSR, AKG, HS, AFG, and HLH

Project administration: AKSR, JFH, LB, AB, PK, NB, PCL, CK, and SMD

Resources: AB, PK, NB, PCL, CK, SMD, NRT, CSS, and MM

Software: AKSR, AKG, HS, AFG, HLH, and LVB

Supervision: AKSR, JFH, NRT, CSS, MM, and DCP

Validation: AKG, HS, DWB, and LVB

Visualization: AKSR, AKG, and HLH

Writing - original draft preparation: AKSR and DCP

Writing - review and editing: AKSR, AKG, DWB, AFG, HLH, LVB, JFH, SD, and DCP

## STAR Methods

### RESOURCE AVAILABILITY

#### Lead contact

Further information and request for resources and reagents should be directed to and will be fulfilled by lead contact, David C. Page.

#### Materials availability

Cell lines are available upon request to the lead contact.

#### Data and code availability

- Raw, de-identified RNA-sequencing data from human cell cultures has been deposited to dbGaP, and processed data has been deposited to github. Accession numbers are listed in the key resources table.
- This paper analyzes existing, publicly available data. Accession numbers for these datasets are listed in the key resources table.
- Original code has been deposited at github and is publicly available as of the date of publication. The repository ID is listed in the key resources table.
- Any additional information required to reanalyze the data reported in this paper is available from the lead contact upon request.

### EXPERIMENTAL MODEL AND SUBJECT DETAILS

#### Human subjects

Adults (18+ years of age) with sex chromosome aneuploidies or euploid controls were recruited through an IRB-approved study at the NIH Clinical Center (12-HG-0181) and Whitehead Institute/MIT (Protocol #1706013503). Informed consent was obtained from all study participants. Individuals with a previous karyotype showing non-mosaic sex chromosome aneuploidy were included in the study. From these individuals, blood samples and skin biopsies were collected at the NIH Clinical Center and shipped to the Page lab for derivation of cell lines. In addition, blood samples from individuals with sex chromosome aneuploidies, and euploid family members, ranging in age from 4-44 years were contributed by the Focus Foundation. Additional LCLs and fibroblast cultures were obtained from the Colorado Children’s Hospital Biobank and Coriell Research Institute, and cultured in the Page laboratory for at least two passages prior to collection for RNA-sequencing. Karyotyping of peripheral blood and fibroblast cell cultures was performed at the National Human Genome Research Institute Cytogenetics and Microscopy Core. To reduce the impact of sex chromosome mosaicism on our sex chromosome aneuploidy analysis, we excluded individuals with >15% mosaicism for other karyotypes. Metadata for cell lines represented in the RNA-sequencing dataset are provided in **Table S1**.

##### Method Details

###### Lymphoblastoid cell lines

Blood was collected in BD Vacutainer ACD tubes and shipped at room temperature to the Page Lab for processing 1-3 d after collection. The buffy coat was resolved by centrifuging blood at 3300 rpm for 10 min, transferred to a new tube with PBS, and subjected to density gradient centrifugation in 50% Percoll (Cytiva) at 3300 rpm for 10 min. Lymphocytes were transferred to a new tube and washed twice with PBS. Lymphocytes were resuspended in 3 mL complete RPMI medium (RPMI 1640 (Gibco), 25mM HEPES (SAFC), 15% FBS (Hyclone), Fungizone (Amphotericin B, Gibco), Gentamicin (Gibco), Penicillin/Streptomycin (Lonza), pH 7.2) per tube of blood and transferred to a T25 flask, supplemented with 0.25mL EBV (produced by B95-8 marmoset lymphoblasts), and 0.2 mL of 1 mg/mL cyclosporine (LC Laboratories). They were incubated for one week at 37°C, fed 1-2 mL complete RPMI, and incubated for another week at 37°C. Once the media began to turn yellow (acidified), cultures were “half-fed” by removing half of the media and replacing it with double the volume. When cultures reached 15 mL, they were transferred to T75 flasks, and gradually expanded to 30 mL, while maintaining a concentration of <1 million cells/mL to ensure viability. Cells were viably frozen for future use by mixing with freezing media (LCL culture media + 5% DMSO), 1 million cells per vial. Cells were also preserved for RNA, DNA, and protein extraction (see below).

###### Primary fibroblast cultures

Our protocol for generating primary skin fibroblast cultures from a skin biopsy is based on (Vangipuram et al., 2013). From adults (18+ years of age) at the NIH Clinical Center we obtained two 4-mm skin punch biopsies from the upper arm, which were immediately placed into a 15 ml conical tube with 10 ml of media (DMEM/F12 (Gibco), 20% FBS, and 100 IU/ml Penicillin-Streptomycin. Tubes were shipped to the Page lab overnight on ice for processing. Each biopsy was used to generate a separate skin fibroblast culture. Biopsies were cut into 18 pieces of equal size and placed 3/well in gelatinized 6-well plates with 1 mL media (High Glucose DMEM (Gibco), 20% FBS, L-Glutamine (MP Biomedicals), MEM Non-Essential Amino Acids (Gibco), 100 IU/ml Penicillin/Streptomycin (Lonza)). Plates were gelatinized by incubating 1 mL sterile 0.1% gelatin (Sigma) solution per well for 30 minutes at room temperature.

Plates were incubated for 1 week at 37°C without disturbance to allow biopsies to attach to the plate and begin to grow out. During week 2, we added 200 μl of fresh media per well every 2-3 days, being careful not to disturb the biopsies. The following week (week 3), we aspirated the media and replaced with 1 mL fresh media per well every 2-3 days. During week 4, we aspirated the media and replaced with 2 mL fresh media per well every 2-3 days. At this point, the fibroblasts generally reached the edges of the wells and were expanded to two T75 gelatinized flasks per 6 well plate. After two days, we combined the cells from the two T75 flasks and split them to three T175 gelatinized flasks. After two days, cells were viably frozen with 1 million cells per vial in freezing media (fibroblast culture media + 5% DMSO). Cells were also preserved for RNA extraction (see below). During optimization of the protocol, cell culture purity was confirmed by immunofluorescence of SERPINH1, a fibroblast marker.

###### Cell collection for subsequent analysis

Cells were collected when LCL cultures reached 30mL, and fibroblasts were ∼80% confluent in three T175 plates. All cell counting was performed using the Countess II cell counter (Life Technologies) and Trypan Blue exclusion. Cultures with >85% cell viability were used in subsequent experiments. To preserve cells for subsequent RNA extraction, 1 million cells were washed in PBS, pelleted, and resuspended in 500 μl TRIzol (Invitrogen) or 200 μl RNAprotect Cell Reagent (Qiagen). Cell suspensions were then frozen at −80°C. Cell cultures were maintained at low passage number; RNA-sequencing experiments were performed on samples at or below passage 4.

Periodically, and on each passage used for experiments, cell cultures were confirmed negative for mycoplasma contamination using either the MycoAlert Kit (Lonza) following the manufacturer’s instructions, or PCR using SapphireAmp Fast PCR Master Mix (Takara) and the following primers:

Myco2(cb): 5’ ctt cwt cga ctt yca gac cca agg cat 3’
Myco11(cb): 5’ aca cca tgg gag ytg gta at 3’

PCR for *GAPDH* was performed on the same sample, using the following primers:

hGAPDH-F: TGT CGC TGT TGA AGT CAG AGG AGA
hGAPDH-R: AGA ACA TCA TCC CTG CCT CTA CTG

Known mycoplasma positive and negative samples were used as a reference.

#### RNA extraction, library preparation, and sequencing

RNA was extracted from 1 million cells per experiment using the RNeasy Plus Mini Kit (Qiagen) following the manufacturer’s instructions, with the following modifications: Cells in RNAprotect Cell Reagent were thawed on ice, pelleted, and lysed in buffer RLT supplemented with 10μl β-mercaptoethanol per mL. For most samples, ERCC control RNAs were added to the lysate based on the number of cells: 10μl of 1:100 dilution of ERCC control RNAs was added per 1 million cells. The lysate was then homogenized using QIAshredder columns (Qiagen), and transferred to a gDNA eliminator column. All subsequent optional steps in the protocol were performed, and RNA was eluted in 30 μl RNase-free water. RNA levels were measured using a Qubit fluorometer and the Qubit RNA HS Assay Kit (ThermoFisher). Before we switched to the per-cell spike-in protocol, we prepared 18 samples in which ERCC control RNAs were added based on amount of RNA after isolation: 2 μl of a 1:100 dilution of ERCC control RNAs was added per 1 μg of RNA. These samples are: #2237, 2245, 6312, 711, 4032, 706, 3429, 3430, 3442, 2690, 2703, 3107, 5297, 5566, 5755, 6029, 2547, and 525. RNA quality control was performed using the 5200 Fragment Analyzer System (Agilent); we consistently purified high-quality RNA with RNA integrity numbers (RIN) near 10. We randomized the samples by karyotype into batches for RNA extraction, library preparation, and sequencing.

RNA sequencing libraries were prepared using the TruSeq RNA Library Preparation Kit v2 (Illumina) with modifications as detailed in (Naqvi et al., 2019), or using the KAPA mRNA HyperPrep Kit V2 (Roche). In both cases, libraries were size selected using the PippinHT system (Sage Science) and 2% agarose gels with a capture window of 300-600 bp. Paired-end 100×100 bp sequencing was performed on a HiSeq 2500 or NovaSeq 6000 (Illumina). **Table S1** lists the library preparation kit and sequencing platform for each sample.

#### RNA-seq data processing and analysis

All analyses were performed using human genome build hg38, and a custom version of the comprehensive GENCODE v24 transcriptome annotation (Godfrey et al., 2020). This annotation represents the union of the “GENCODE Basic” annotation *and* transcripts recognized by the Consensus Coding Sequence project (Pruitt et al., 2009). Importantly, the GENCODE annotation lists the PAR gene annotations twice – once on Chr X and once on Chr Y– which complicates analysis. We removed these annotations from Chr Y so the PAR genes are only listed once in our annotation, on Chr X. To analyze samples in which ERCC spike-ins were added, we merged our custom transcript annotation with the ERCC Control annotation.

Reads were pseudoaligned to the transcriptome annotation, and expression levels of each transcript were estimated using kallisto software v0.42.5 (Bray et al., 2016). We included the “--bias” flag to correct for sequence bias. The resulting count data (abundance.tsv file) were imported into R with the tximport package v1.14.0 (Soneson et al., 2015) for normalization using DESeq2 v1.26.0 (Love et al., 2014). For downstream analysis, we used only protein-coding genes (as annotated in ensembl v104) with the following exceptions: we included genes annotated as pseudogenes on Chr Y that are members of X-Y pairs (*TXLNGY, PRKY*) and well-characterized long non-coding RNAs (lncRNAs) involved in X-inactivation or other processes (*XIST, JPX, FTX, XACT, FIRRE, TSIX).* We annotated genes distal to *XG*, which spans the pseudoautosomal boundary on Xp and is truncated on Chr Y, as part of PAR1 - 15 genes in total. PAR2 comprised the four most distal genes on Xq and Yq. Annotations of non-pseudoautosomal region of the X (NPX) genes with homologs on the non-pseudoautosomal region of the Y (NPY) were derived from (Bellott and Page, 2021). 224 protein-coding genes on Chr 21 (ensembl v104) were used as a starting point for our analyses. We excluded 21 annotated genes in several regions with high homology between the long and short arms of Chr 21 because the assembly was not fully validated in these regions (https://www.ncbi.nlm.nih.gov/grc/human/issues?filters=chr:21).

#### Identifying genes affected by changes in Chr X, Y, or 21 copy number

We first defined lists of expressed NPX, NPY, PAR, or Chr 21 genes as those with median TPM of at least 1 in 46,XX or 46,XY samples. To ensure that no genes with robust expression were excluded, we also analyzed LCL and fibroblast expression data from GTEx (GTEx Consortium, 2017), and included several genes that were just below our TPM cutoff but had median TPM of at least 1 in those datasets.

For each expressed NPX, NPY, or PAR gene we performed linear modeling using the lm() function in R. These calculations suppose that each additional chromosome adds a consistent and equal increment to the total expression level of the gene in question.

For NPX and PAR genes we used the following equation:

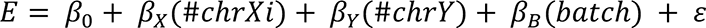

E represents the expression (read counts) per gene, 𝛽_0_ represents the intercept, 𝛽_𝑋_ and 𝛽_𝑌_ are the coefficients of the effect of additional copies of Chr Xi or Y, respectively, and ∈ is an error term. For this equation, the intercept represents the 45,X samples.

For NPY genes we employed the following equation, analyzing only those samples with one or more copies of Chr Y:

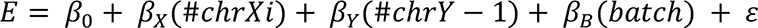

For this equation, the intercept represents the 46,XY samples.

For Chr 21 genes we employed the following equation, analyzing only those samples with 46,XX; 46,XX; 47,XY,+21; or 47,XX,+21 karyotypes:

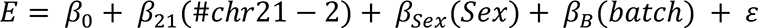

𝛽_21_and 𝛽*_sex_* are the coefficients of the effect of an additional copy of Chr 21 and sex (XY vs XX), respectively. For this equation, the intercept represents the 46,XX samples.

The resulting *P*-values were adjusted for multiple hypothesis testing using the p.adjust() function in R, specifying the Benjamini Hochberg method. Genes with a false discovery rate (FDR) < 0.05 were considered significant. To compute the normalized expression change per Chr Xi (ΔE_X_) or Y (ΔE_Y_), we divided the coefficient of interest (𝛽_X_ or 𝛽_Y_) by the average intercept across batches, which corresponds to the baseline expression of the gene in samples with only one X chromosome (for NPX and PAR genes) or one Y chromosome (in the case of NPY genes). For Chr 21, we computed ΔE21 by dividing the coefficient (𝛽_21_) by the average intercept across batches divided by two to obtain the average expression from one copy of Chr 21.

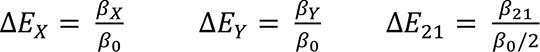

In the case of *XIST*, which is only expressed when two or more copies of Chr X are present, we used the following equations:

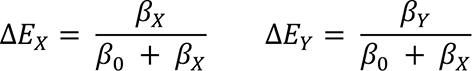

We calculated the standard error of ΔE_X_, ΔE_Y_, and ΔE_21_ using the following equations:

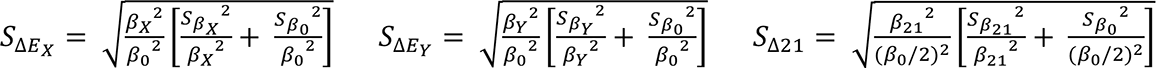

To confirm the validity of our approach, we used bootstrapping to sample our dataset with replacement 1000 times and obtained similar results. *BEX1* was removed from downstream analyses in fibroblasts because two samples (one 45,X and one 49,XXXXY) had high expression values for this gene resulting in > 25 times higher error values for ΔE_X_ and ΔE_Y_ compared to all other genes.

#### Saturation analysis for sex chromosome-encoded genes

For LCLs and fibroblasts, size-*n* subsets of available RNA-seq libraries were sampled randomly without replacement, 100 times for each sample size, *n*. After confirming that the model matrix would be full rank in each sampling (for example, that samples would not all be of the same karyotype or batch), we performed linear modeling on NPX, PAR, NPY, and Chr 21 genes as described above to identify genes whose expression changes significantly (FDR < 0.05) with copy number of Chr X, Y or 21.

#### Assessing linearity of sex-chromosome gene expression changes

To assess whether sex-chromosome gene expression changed linearly (*i.e.*, by a fixed amount) with additional X or Y chromosomes, their expression levels across the LCL or fibroblast samples were fit by non-linear least squares to the power curves shown below, using the “nlsLM” function from the R package “minpack.lm”.

NPX genes:

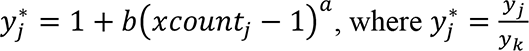

PAR genes:

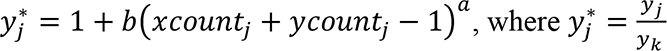

NPY genes:

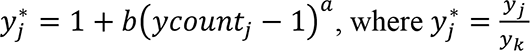

In each of the equations above, 𝑦_𝑗_^∗^ is the normalized RNA-seq read count for a given gene in sample 𝑗, given by the raw read count in sample 𝑗 divided by the average read count in the set of samples 𝑆_{·}_ with only one chromosome of the relevant type: for NPX genes, 1 copy of Chr X (and any number of Y chromosomes); for PAR genes, 45,X samples; for NPY genes, 1 copy of Chr Y (and any number of X chromosomes). 𝑏 = 0.5 and 𝑎 = 1 were used as initial parameter values. Fitted values of 𝑎 ≈ 1 indicate a linear relationship between expression and sex-chromosome count. Fitted values of 𝑎 ≈ 0 or 𝑏 ≈ 0 indicate no change in expression with X or Y count.

#### 𝛥E_X_ calculations in samples with 0 Y chromosomes (females) and 1 Y chromosome (males)

We took subsets of the samples with either zero Y chromosomes (females) and with one Y chromosome (males) and performed the same linear modeling and 𝛥E_X_ calculations as above. We removed *MAP7D2* in female LCLs, *IL13RA2* in female fibroblasts, and *FHL1* in male fibroblasts because their error values (likely due to smaller sample size) were much higher than those of other genes. To compare the linear modeling results, we performed Pearson correlations between the results using all samples, and those from male-only or female-only samples.

#### Reanalysis of array data and comparison to RNA-seq data

A previous study performed gene expression analysis, using Illumina oligonucleotide BeadArrays, of LCLs from 68 individuals of the following karyotypes: 45,X; 46,XX; 46,XY; 47,XXX; 47,XXY; 47,XYY; and 48,XXYY (Raznahan *et al*., 2018). Since this microarray dataset was generated from an independent set of samples, we sought to validate our results through a reanalysis of the data.

The raw data from the microarrays was not publicly available, but the authors provided us pre-processed data upon request, which we used to perform our analysis. To identify genes that cleared a minimum signal threshold to be considered expressed in the microarray data, we assessed the median signal in 46,XY samples for all Chr Y genes annotated on the microarray. We focused on Chr Y genes in this analysis because many are known to be expressed exclusively in testes, and therefore could provide us with an appropriate sense of the background signal expected for genes not expressed in LCLs. From this analysis, we concluded that a signal threshold of 111 would be appropriate for identifying expressed genes. We used this threshold to identify 278 expressed Chr X genes (including PAR and NPX genes) in the microarray dataset. This was fewer than the 341 expressed Chr X genes identified in our LCL RNA-seq data, but more than double the 121 expressed Chr X genes considered in Raznahan *et al*. (Supplemental Table 4 in (Raznahan *et al*., 2018)). This discrepancy could not be resolved by simply increasing the signal threshold in our analysis, as *TMSB4X,* one of the most highly expressed genes in LCLs, was excluded from Raznahan and colleagues’ list of expressed genes.

Using our list of 278 expressed genes from the Raznahan *et al*. dataset, we analyzed the microarray signal values (in place of RNA-seq read counts) using linear models as a function of Xi copy number, controlling for Chr Y copy number. We calculated 𝛥E_X_ values from the microarray data and compared these to our RNA-seq dataset using a Pearson correlation, which revealed that the results were generally concordant. For genes that were lowly expressed, however, 𝛥E_X_ values tended to be much lower in the microarray dataset, consistent with the higher sensitivity of RNA-seq data.

#### Isoform-specific analysis of RNA-seq data

After estimating counts for each transcript using kallisto software (as described above, with 100 bootstraps), we used sleuth v0.30.0 to normalize those transcript counts (Pimentel et al., 2017). X chromosome transcripts were called as expressed if their corresponding gene was on the list of expressed genes (above) and median transcript counts were > 200. Linear regressions and 𝛥E_X_ calculations were performed as for genes (above) to identify transcripts whose abundance changes significantly with additional copies of Chr X.

The following ENCODE datasets were used for visualization in IGV software (Robinson et al., 2011) at the *UBA1* locus:

| Assay | Cell type | Cell line | Karyotype | Accession |
| --- | --- | --- | --- | --- |
| CTCF<br>ChIP-seq | LCL | GM12878 | 46,XX | ENCFF644EEX |
|  |  | GM12864 | 46,XY | ENCFF070FTG |
|  | Fibroblast | AG09309 | 46,XX | ENCFF233THH |
|  |  | AG10803 | 46,XY | ENCFF080HIA |
| CTCF<br>ChIA-PET | LCL | GM12878 | 46,XX | ENCFF80PGS<br>ENCFF847QOE |

#### Gene constraint analysis

To investigate sensitivity to a reduction in gene dosage, we used three metrics: LOEUF, RVIS, and pHI. We downloaded LOEUF (loss-of-function observed/expected upper fraction) scores from gnomAD (v2.1.1.lof_metris.by_gene.txt; https://gnomad.broadinstitute.org/), and only used scores with a minimum of 10 expected LoF variants. Updated RVIS (residual variation intolerance scores) from (Petrovski *et al*., 2013) including the ExAC dataset were downloaded from http://genic-intolerance.org/data/RVIS_Unpublished_ExAC_May2015.txt. Updated probability of haploinsufficiency (pHI) scores from (Huang *et al*., 2010) were downloaded from https://www.deciphergenomics.org/files/downloads/HI_Predictions_Version3.bed.gz. To complement these data, we obtained a list of genes with observed homozygous loss-of-function variants from (Karczewski *et al*., 2020). For sensitivity to an increase in gene dosage, we used the per-gene average probability of conserved miRNA targeting scores (P_CT_) (Friedman *et al*., 2009).

For each metric, we computed a percentile rank score, ranking from most-to least-constrained. Because several of the metrics calculated scores separately for autosomal (including PAR genes) and NPX genes, we ranked autosomal (and PAR) genes separately from NPX genes. All annotated genes, regardless of expression status in LCLs or fibroblasts, were included in the rankings, with the following exceptions: 1) NPX genes previously annotated as “ampliconic” in (Jackson et al., 2021; Mueller et al., 2013), since constraint metrics cannot be accurately applied to these highly similar genes, and 2) genes with <2 annotations across all metrics.

To obtain an aggregate sense of a gene’s expression constraint across multiple metrics, we calculated the average. Among NPX genes, we considered those with |ΔE_X_ ≥ 0.1| (FDR<0.05) to be most likely to contribute to phenotypes mediated by Xi copy number, prioritizing the top ten genes by the average gene-constraint metric. For PAR1 genes, we prioritized genes with an average gene constraint percentile ranking of at least 50%. To assess the phenotypic roles of highly-constrained genes, we annotated them for disease phenotypes with known molecular basis from Online Mendelian Inheritance in Man (OMIM) (McKusick-Nathans Institute of Genetic Medicine, 2022).

#### Comparisons to published annotations of X-inactivation status

We re-compiled XCI status annotations of individual genes from four studies of Chr X allelic ratios: (Cotton *et al*., 2013; Garieri *et al*., 2018; Sauteraud *et al*., 2021; Tukiainen *et al*., 2017). Previous XCI status compilations (Balaton *et al*., 2015) incorporated DNA methylation data, which we excluded because it does not directly measure Xi transcription. Previous compilations also incorporated information about expression in human-rodent hybrid cell lines carrying a human Xi (Carrel and Willard, 2005); we incorporated this information only where allelic ratios were not available. Our final XCI status annotations are listed in **Table S6A**, with the workflow for generating these annotations explained below.

The first dataset that we incorporated was derived from paired genomic and cDNA SNP-chips in skewed LCL and fibroblast cell cultures (Additional file 7 in (Cotton *et al*., 2013)). We used the AR values provided (average Xi expression column) for genes informative in at least 5 samples, resulting in AR values for 424 genes. Using the provided numbers of informative samples and standard deviations of AR values, we computed 95% confidence intervals for the AR values. We considered a gene “Subject” to XCI if the AR 95% confidence interval included zero or the AR value was <0.1; otherwise we considered the gene to “Escape”.

The second dataset that we incorporated was derived from bulk or single cell RNA-seq of LCLs (Tukiainen *et al*., 2017). The bulk RNA-seq was from an individual in the GTEx dataset with 100% skewed XCI across the body (Table S5 in (Tukiainen *et al*., 2017)). The single-cell RNA-seq in LCLs was from three individuals (Table S8 in Tukiainen *et al*.; we excluded data from one dendritic cell sample). For each dataset, we calculated an AR for each gene using read counts from the more lowly and highly expressed alleles in each sample, and used the provided adjusted P-values to identify genes with significant Xi expression (padj < 0.05). For a gene to be considered informative, we required data from at least two individuals in the single cell dataset, or one individual in the single cell dataset and informative in the bulk RNA-seq dataset, resulting in 82 informative genes. We called a gene as “Subject” to XCI if there was no significant expression from Xi in either the bulk or single-cell datasets, and “Escape” if one or both of the datasets showed evidence of Xi expression.

The third dataset that we incorporated was derived from single-cell allelic expression in fibroblasts (Garieri *et al*., 2018). The dataset includes five individuals (Dataset 3 in Garieri *et al*.), and we required data from at least two samples to be considered informative for a given gene, resulting in 203 genes. We converted their reported values (Xa reads/total reads) to AR values using the following formula: 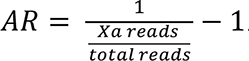 We used the AR threshold calculated in (Garieri *et al*., 2018) to consider a gene significantly expressed from Xi in each sample (AR>0.0526). If a gene had no samples with significant expression from Xi or a mean AR value <0.1 across samples, we considered it “Subject” to XCI; otherwise, it was judged to “Escape” XCI.

The fourth dataset that we incorporated was derived from allele-specific bulk RNA-seq performed on 136 samples with skewed XCI from the set of GEUVADIS LCLs (Tables S4 and S5 in (Sauteraud *et al*., 2021)). For a gene to be considered scorable, we required that it be informative in at least 10 samples, resulting in 215 genes. We calculated an AR for each gene in each sample using the read counts from the more lowly and highly expressed alleles in each sample, adjusting for the level of skewing in each sample. To identify genes that were significantly expressed from Xi across samples, we performed paired, two-sample, one-sided t-tests using the t.test function in R, asking whether the raw (pre-adjusted for skewing) AR values were greater than the baseline AR given the level of skewing in each sample 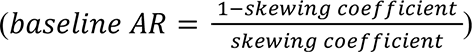; we corrected the resulting P-values for multiple comparisons with the p.adjust function in R using the Benjamini-Hochberg method. Genes were considered “Escape” if they had padj < 0.01, and “Subject” otherwise.

Next, we synthesized the calls from these four datasets. We assigned a gene as “Subject” if: 1) all studies were “Subject” or 2) most (>50%) studies were “Subject” and the average AR across all studies was <0.1. We assigned “Escape” if 1) most (>50%) studies were “Escape” or 2) 50% or fewer (but more than 0) studies were “Escape” and either i) there was more than one study with evidence of escape or ii) the average AR across all studies was <0.1. Finally, we assigned “No call” if the gene was not informative in any of the four datasets. For these genes, we investigated whether there were any calls using hybrids from (Carrel and Willard, 2005) as compiled in (Balaton *et al*., 2015). If a gene had no call in any of the four AR datasets, but had a proportion of expression in Xi hybrids <0.22, we considered the gene “Subject”; genes with a greater proportion were called “Escape.”

To compare our calls with previous XCI consensus calls, we made the following modifications to the Balaton list: *XG* had been listed as a PAR gene, but we excluded it from our list of PAR genes because it is located at the PAR boundary and truncated on the Y chromosome. We updated its annotation to escape (“E”) since the Balaton table lists evidence for escape. The Balaton table lists *XIST* as “mostly subject” to XCI, but given its exclusive expression from Xi, we updated its status to escape (“E”). We manually examined all genes on our list that were not found in the Balaton list to make sure that genes were not misclassified due to differences in official gene names. For those genes still not present in the Balaton list after this correction, we list “No call”. To compare with our annotations, we grouped the Balaton calls into “Escape” if they were annotated as “PAR”, “Escape”, “Mostly escape”, “Variable Escape”, “Mostly Variable Escape”, or “Discordant”. We grouped Balaton calls into “Subject” if they were annotated as “Mostly subject” or “Subject”.

We compared our new calls with the Balaton calls for the 423 genes expressed in fibroblasts or LCLs, finding 48 where they differed. Of these, nine had a call in Balaton, but no call in the newer datasets. For two genes (*TCEAL3*, *TMSB4X*), there was no call in Balaton, but newer data enabled a call to be made (both “Subject”). Nineteen genes were called as “Subject” in Balaton, but new data indicates that they have expression from Xi and we categorize them as “Escape.” The final eighteen genes were called as “Escape” in Balaton, but new data suggested they have no expression from Xi. In total, our classification found 86 genes that “Escape”, 315 genes that are ”Subject” to XCI, and 22 genes with “No call.”

#### Allele-specific expression analysis

##### SNP calling

We called SNPs in each RNA-seq sample with two X chromosomes (46,XX, 46,XX,+21, 47,XXY, 48,XXYY) following the Broad Institute’s “Best Practices” workflow for identifying short variants in RNA-seq data (https://gatk.broadinstitute.org/hc/en-us/articles/360035531192-RNAseq-short-variant-discovery-SNPs-Indels-). To perform our skewing analysis, we filtered for SNPs with the following properties: 1) annotated in the dbSNP database, 2) located in an exon of an expressed gene, 3) displaying a minimum coverage of 10 reads, and 4) heterozygous with at least three reads mapping to each of the reference and alternative alleles. We excluded SNPs where the presence of two alleles likely represented technical artifacts rather than biallelic expression, including in *WASH6P* (SNPs map to multiple near-identical autosomal paralogs), *ATRX* (SNP in a mutation-prone stretch of Ts), and *APOOL* (SNPs within an inverted repeat).

For samples with a copy of Chr Y, we excluded SNPs mapping to PAR genes, to avoid measuring allelic contributions of Chr Y.

##### Identifying cell lines with skewed X chromosome inactivation

We classified genes as “Xa-only” (only expressed from Xa) if previously characterized as “silenced” *and* found here to have ΔE_X_<0.05 (FDR>0.5); see **Table S6E**. We expect that in skewed cell lines, reads from Xa-only genes should be near or completely monoallelic. For each SNP in Xa-only genes, we calculated the “skewing coefficient” by dividing the number of reads from the dominant allele by the total number of reads covering the SNP. These coefficients range from 0.5 (equal expression of two alleles) to 1 (expression from a single allele). For each sample, we computed the median skewing coefficient across all SNPs in Xa-only genes, requiring a threshold of 0.8 to classify as skewed. Using simulations, we find that this level of skewing is unlikely to occur by chance (P<1×10^-6^), and we do not find evidence of such skewing for SNPs on Chr 8, an autosome with a similar number of expressed genes.

Several samples had few (≤ 5) informative SNPs in Xa-only genes, but many SNPs in other genes. We interpret this to mean that these samples are highly skewed and that we do not observe enough RNA reads covering both alleles to count SNPs in Xa-only genes as informative. Between these highly-skewed samples and the samples with skewing coefficients of at least 0.8, we identified 21 LCLs and 10 fibroblast cultures with skewed XCI.

##### Determining allelic ratios for X chromosome genes

After identifying the skewed cell lines, we identified genes with informative SNPs values in at least three skewed samples of a given cell type. We then computed the allelic ratio (AR) at each informative SNP by dividing the number of reads from the more lowly expressed allele by the number of reads from the more highly expressed allele. In cell cultures that are partially skewed, genes will appear more biallelic than in completely skewed cell cultures since there are two populations of cells with different active X chromosomes present – the “major” and “minor” cell populations. Using our skewing estimates, we adjusted the AR on a per-sample basis using the following formula:

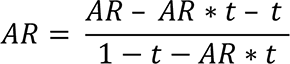

Where t is the estimated percentage of cells in the “minor” population (*i.e.*, with the other X chromosome active compared to the “major” cell population), calculated by: 1 – skewing coefficient. For highly-skewed samples, we were unable to calculate a stringent skewing coefficient due to too few SNPs, so we set skewing coefficient = 1. As a result, it is possible that allelic ratios in these samples may be slightly overestimated if the skewing coefficients are in fact <1. Within each sample, we obtained the average AR for each gene by averaging across all informative SNPs in that gene’s exons and then calculated the mean AR across skewed samples to obtain a final per-gene AR estimate.

To assess whether AR values for each gene were significantly greater than zero, we performed one-sided t-tests using the t.test function in R, asking whether the AR values were greater than zero; we corrected the resulting P-values for multiple comparisons with the p.adjust function in R using the Benjamini-Hochberg method. We also repeated this analysis excluding highly skewed samples, since the skewing coefficients cannot be stringently determined. This removed some informative genes but did not significantly affect the AR values (**Note S2**).

To identify genes whose AR and 𝛥E_X_ values differ significantly, we performed one-sample, two-sided t-tests for the AR values across samples, setting mu= 𝛥E_X_. We selected genes with Benjamini-Hochberg adjusted P-values <0.1 as having significantly different AR and 𝛥E_X_ values. From this list we excluded genes for which the 95% confidence interval of 𝛥E_X_ values (1.96*standard error) included the mean AR value, and those for which both 𝛥E_X_ and AR were not significantly different from zero (FDR≥ 0.05).

We compared our AR values derived from LCLs or fibroblasts with the four published allelic-ratio datasets described in the above methods on generating XCI status calls (**Note S2**).

### QUANTIFICATION AND STATISTICAL ANALYSES

Various statistical tests were used to calculate P-values as indicated in the methods section, figure legend, or text, where appropriate. Results were considered statistically significant when P<0.05 or FDR<0.05 when multiple hypothesis correction was applied, unless stated otherwise. Data are shown as median and interquartile range, unless stated otherwise. All statistics were calculated using R software, version 3.6.3 (R Development Core Team, 2020).

### KEY RESOURCES TABLE

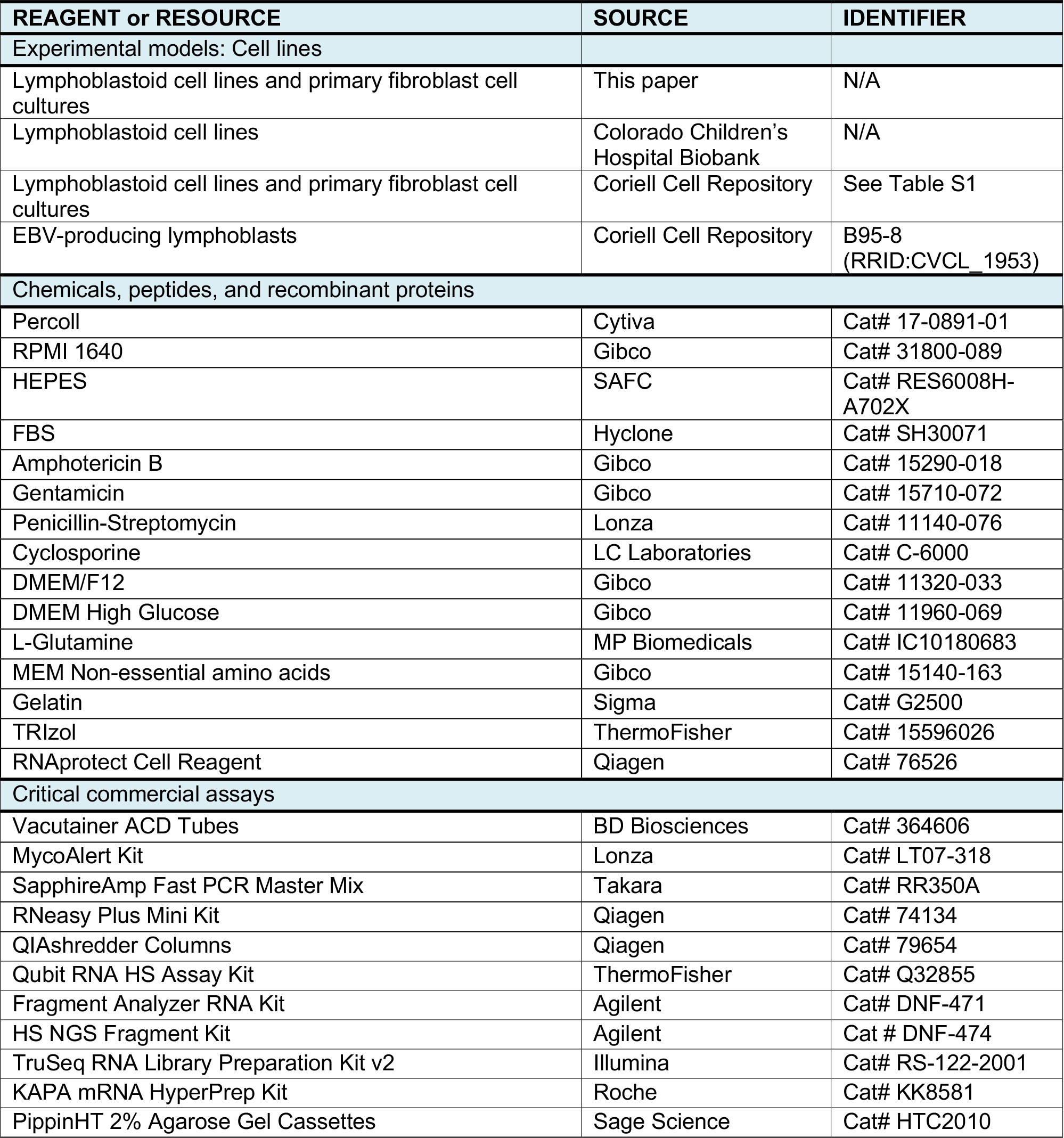
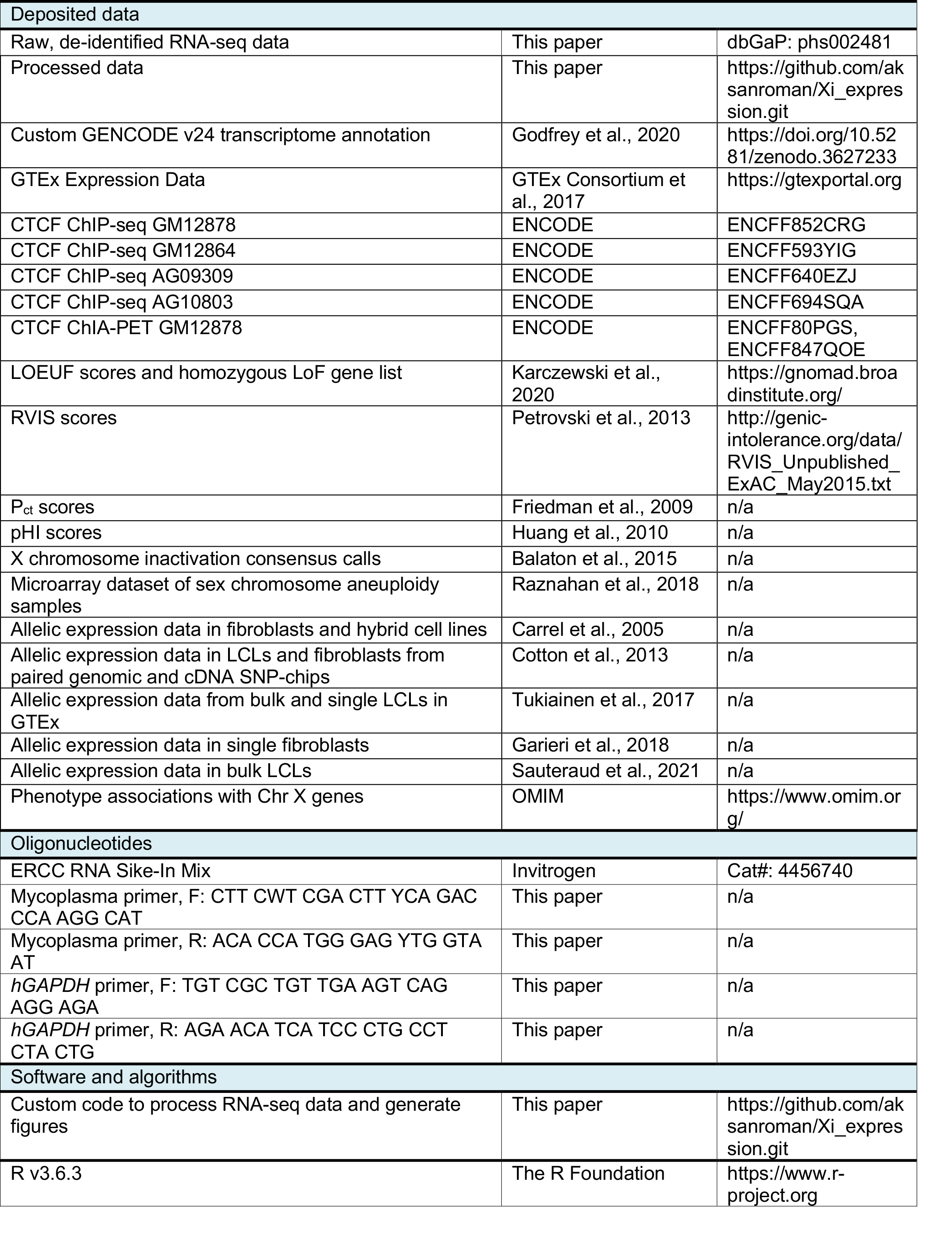
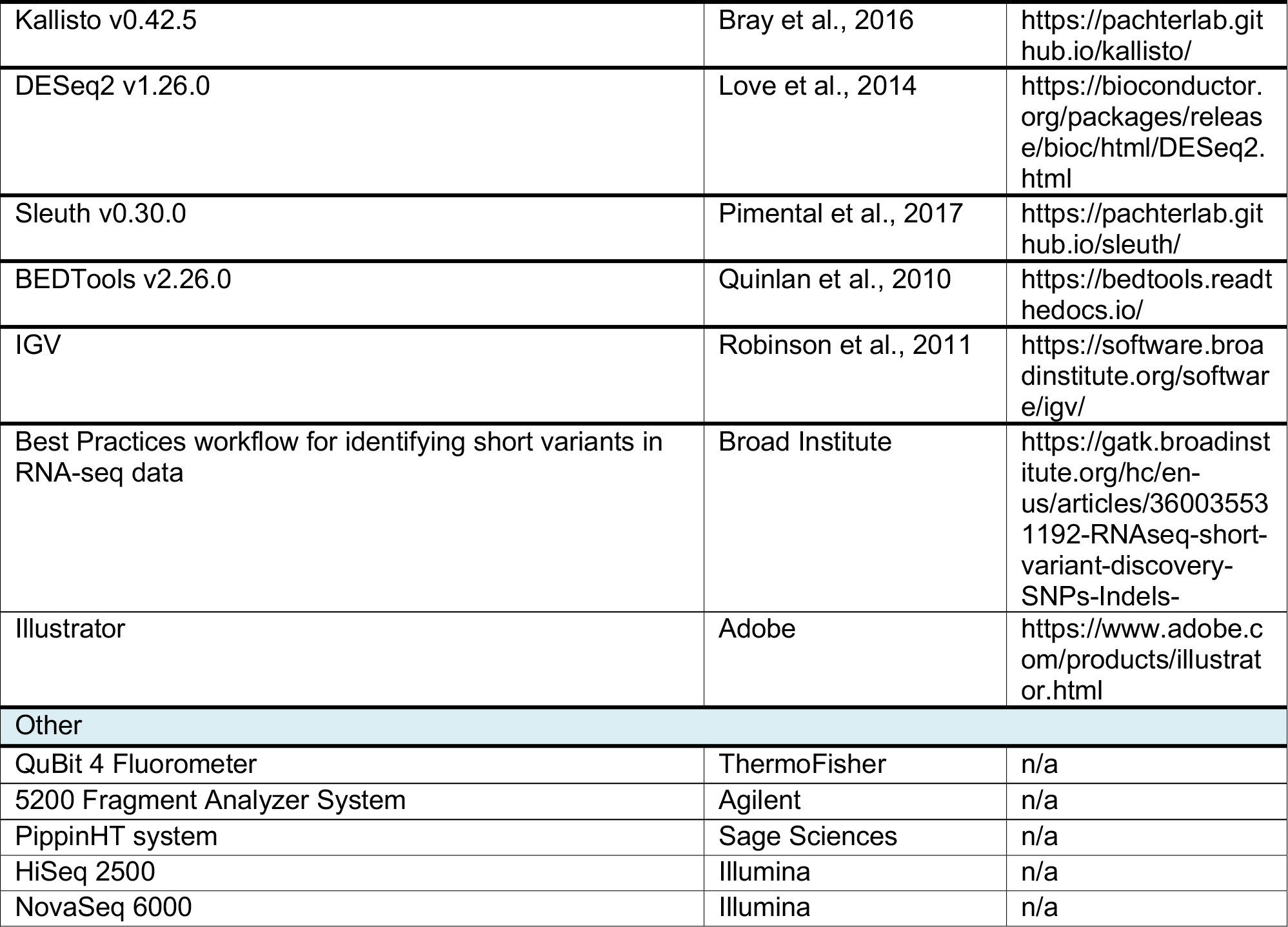

## Supplemental information

**Figure S1. Bootstrapping analyses reveal that few additional significant genes would be observed with a larger sample size.** Each analysis was performed by randomly choosing the indicated number of samples and performing the linear regression analysis, repeated 100 times for each sample size. The magenta point in each figure represents the number of significant genes in the final analysis using all of the samples. **(A)** NPX genes with Y homologs with ΔE_X_>0 reach saturation by a sample size of 40. (**B)** For NPX genes without Y homologs, the number of genes with ΔE_X_>0 increases rapidly at low sample sizes, and then levels off. **(C)** For NPX genes with ΔE_X_<0, more samples are required to see an initial increase in the number of significant genes. In LCLs, 50-60 samples are needed to identify 20 genes with ΔE_X_<0, whereas only 30-40 samples are needed to identify 20 genes with ΔE_X_>0. For LCLs, the maximum increase was observed between 60-70 samples where a median of 7 genes were added, with subsequent increases being less (5 genes from 70-80, 3.5 from 80-90 and 3.5 from 90-100). In fibroblasts, the maximum increase was observed between 40-50 samples where a median of 4.5 genes were added, with only 3 or 4 additional genes with each additional 10 samples. If this trajectory continues, these results indicate that additional samples would only result in a few additional genes being identified as significant. **(D)** PAR genes with both ΔE_X_>0 and ΔE_Y_>0 reach saturation by sample size of 50. **(E)** NPY genes with ΔE_Y_>0 reach saturation after 25 samples. **(F)** The trajectory for Chr 21 genes with ΔE_21_>0 increases rapidly between 10-30 samples, and then slows.

**Figure S2. Linear functions are a good fit for modeling sex chromosome aneuploidy RNA-seq data.** For sex chromosome genes in LCLs and fibroblasts, power functions were fit (gray boxes) by non-linear least squares regression. Scatterplots show fitted values for the exponent parameter, *a*, and the coefficient, *b*, for each gene. *XIST* and genes for which the regression model did not converge are excluded from the plots. **(A-B)** Some NPX genes cluster near *a*=1, indicating their expression increases by a fixed amount for each additional X or Y chromosome. These correspond to genes previously annotated as escaping XCI **(C-D)**. Other NPX genes cluster near *a*=0 or *b*=0, indicating that they do not change in expression with additional copies of Xi, and correspond to genes previously annotated as subject to XCI (**E-F**). (**G-H**) PAR1 genes cluster near a=1, indicating their expression increases by a fixed amount for each additional sex chromosome, in contrast to PAR2 genes. (**I-J**) NPY genes also cluster near a=1, indicating their expression increases by a fixed amount for each Y chromosome.

**Figure S3. Extended examples of** Δ**E values for Chr X, Y and 21 genes.** Scatterplots and regression lines with confidence intervals for selected genes across Chr X, Y, or 21 copy number series. Adjusted p-values (FDR) <0.05 indicate that ΔE_X_ values are significantly different from zero. **(A)** Fifteen Chr X genes with ΔE_X_=0 in LCLs and fibroblasts. **(B)** Fifteen Chr X genes with ΔE_X_>0 in LCLs and fibroblasts. **(C)** Seven Chr X genes with ΔE_X_<0 in LCLs and fibroblasts. **(D)** All expressed Chr Y genes. **(E)** Fifteen Chr 21 genes with ΔE_21_>0.

**Figure S4.** Δ**E_X_ values are similar when calculated with all samples, or with female (0 Y chromosomes) or male (1 Y chromosome) samples.** Scatterplots of all expressed genes in LCLs (A-B) or fibroblasts (C-D) showing ΔE_X_ values calculated using all samples (X-axes) or exclusively female samples (A, C Y-axes) or male samples (B, D Y-axes). Deming regression lines, Pearson correlations and P-values are indicated.

**Figure S5. Quantitative analysis of Xi contributions to transcript isoform expression (A)** Accounting of Chr X genes with multiple expressed transcript isoforms in LCLs and fibroblasts. Indicated here are the numbers of genes where i) all isoforms show ΔE_X_≈0, or ii) all isoforms show ΔE_X_ values significantly different from zero in the same direction (FDR<0.05), or iii) isoforms show discordant ΔE_X_ values. **(B)** Venn diagram shows the overlap of genes expressed in both LCL and fibroblasts that had isoforms with discordant ΔE_X_ values. **(C)** Scatterplots of each transcript isoform’s ΔE_X_, compared with the gene’s ΔE_X_ in LCLs and fibroblasts. Three *ZFX* isoforms are indicated with blue points; two *UBA1* isoforms indicated with red points. Pearson correlations (r) between isoforms and genes and P-values are indicated. **(D)** Scatterplots of expression of three *ZFX* isoforms across Chr X copy number in LCLs and fibroblasts showing that they consistently have ΔE_X_>0. **(E)** Scatterplots of expression of two *UBA1* isoforms across Chr X copy number in LCLs and fibroblasts showing discordant ΔE_X_ values. Regression lines, ΔE_X_, and FDR are indicated.

**Figure S6. Reanalysis of microarray data from LCLs of individuals with varying sex chromosome constitutions using linear modeling. (A)** Chr Y gene expression in microarray data from XY samples was assessed to set a minimum signal threshold of 111 (red line) for expressed genes, based on those expressed in LCL RNA-seq. **(B)** ΔE_X_ values in RNA-seq and microarray data for expressed Chr X genes were highly correlated. Deming regression line, Pearson correlation and P-value are indicated.

**Note S1.** Divergent ΔE_X_ values for transcript isoforms of an X-linked gene

**Note S2.** Identifying cell lines with skewed X chromosome inactivation from RNA-seq data

**Table S1.** Metadata for euploid and aneuploid samples with RNA-sequencing

**Table S2.** Linear regression results for expressed PAR and NPX genes

**Table S3.** Linear regression results for expressed PAR and NPX transcript isoforms

**Table S4.** Linear regression results for expressed NPY genes

**Table S5.** Linear regression results for expressed Chr 21 genes

**Table S6.** Allelic ratios and statistics for informative Chr X genes

**Table S7.** Expression constraint metrics for PAR1 and NPX genes.

