## Supplemental Information for "The human inactive X chromosome modulates expression of the active X chromosome"

**Figures S1-S6**

**Note S1-S2**

**Tables S1-S7** *(provided as separate .xlsx files)*

**Figure S1**

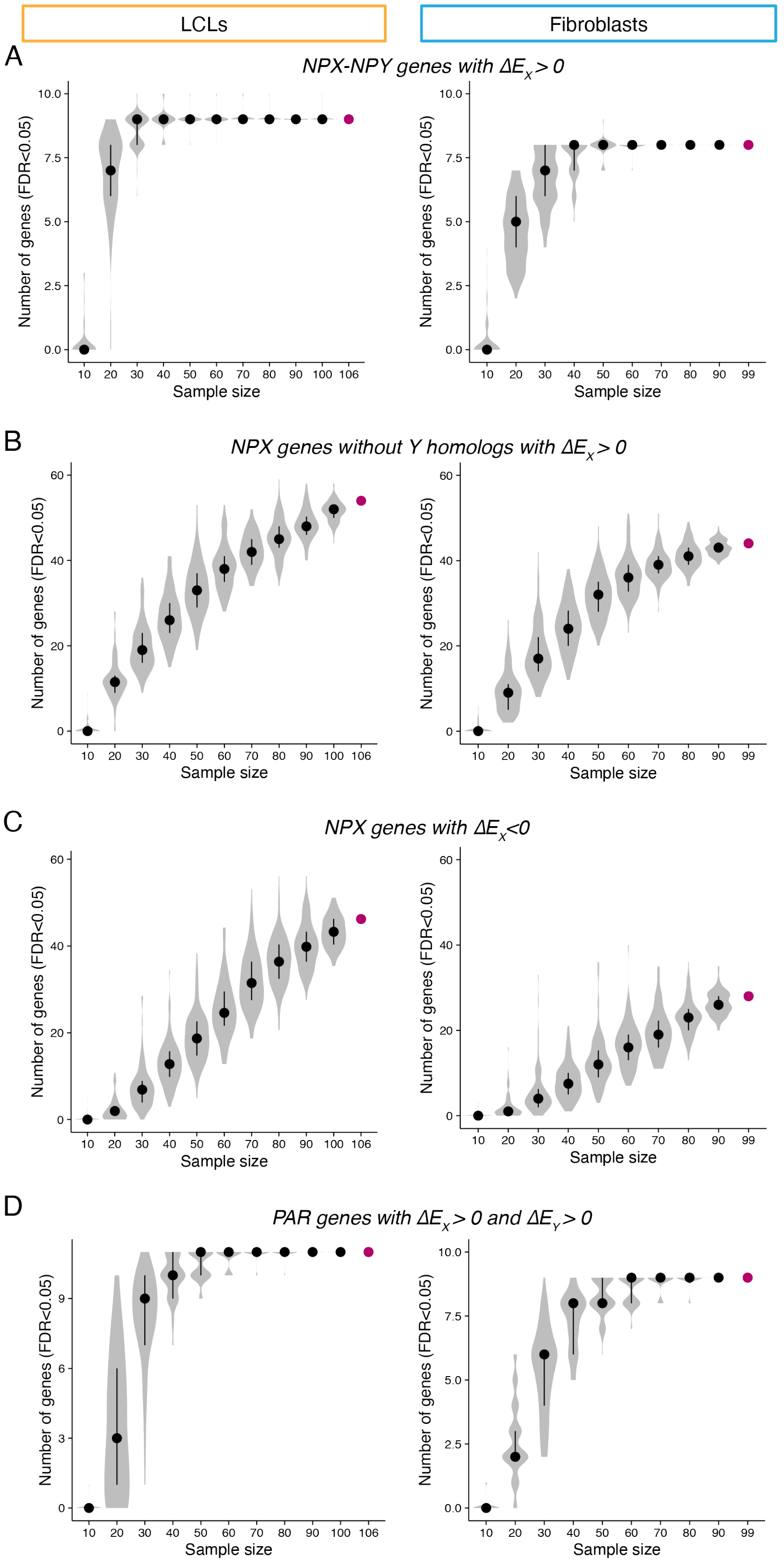
LCLs
Fibroblasts
A
NPX-NPY genes with ΔE$_X$ > 0
B
NPX genes without Y homologs with ΔE$_X$ > 0
C
NPX genes with ΔE$_X$<0
D
PAR genes with ΔE$_X$ > 0 and ΔE$_Y$ > 0
Number of genes (FDR<0.05)
Sample size

**Figure S1, continued.**

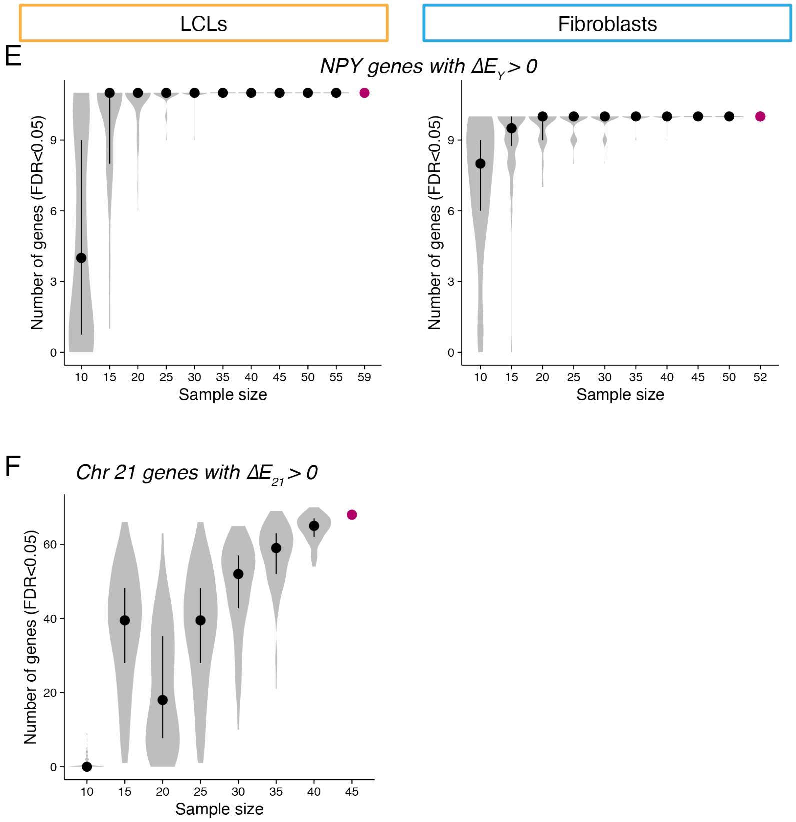


**Figure S1. Bootstrapping analyses reveal that few additional significant genes would be observed with a larger sample size.** Each analysis was performed by randomly choosing the indicated number of samples and performing the linear regression analysis, repeated 100 times for each sample size. The magenta point in each figure represents the number of significant genes in the final analysis using all of the samples. **(A)** NPX genes with Y homologs with $\Delta E_X>0$ reach saturation by a sample size of 40. (**B)** For NPX genes without Y homologs, the number of genes with $\Delta E_X>0$ increases rapidly at low sample sizes, and then levels off. **(C)** For NPX genes with $\Delta E_X<0$, more samples are required to see an initial increase in the number of significant genes. In LCLs, 50-60 samples are needed to identify 20 genes with $\Delta E_X<0$, whereas only 30-40 samples are needed to identify 20 genes with $\Delta E_X>0$. For LCLs, the maximum increase was observed between 60-70 samples where a median of 7 genes were added, with subsequent increases being less (5 genes from 70-80, 3.5 from 80-90 and 3.5 from 90-100). In fibroblasts, the maximum increase was observed between 40-50 samples where a median of 4.5 genes were added, with only 3 or 4 additional genes with each additional 10 samples. If this trajectory continues, these results indicate that additional samples would only result in a few additional genes being identified as significant. **(D)** PAR genes with both $\Delta E_X>0$ and $\Delta E_Y>0$ reach saturation by sample size of 50. **(E)** NPY genes with $\Delta E_Y>0$ reach saturation after 25 samples. **(F)** The trajectory for Chr 21 genes with $\Delta E_{21}>0$ increases rapidly between 10-30 samples, and then slows.

**Figure S2.**

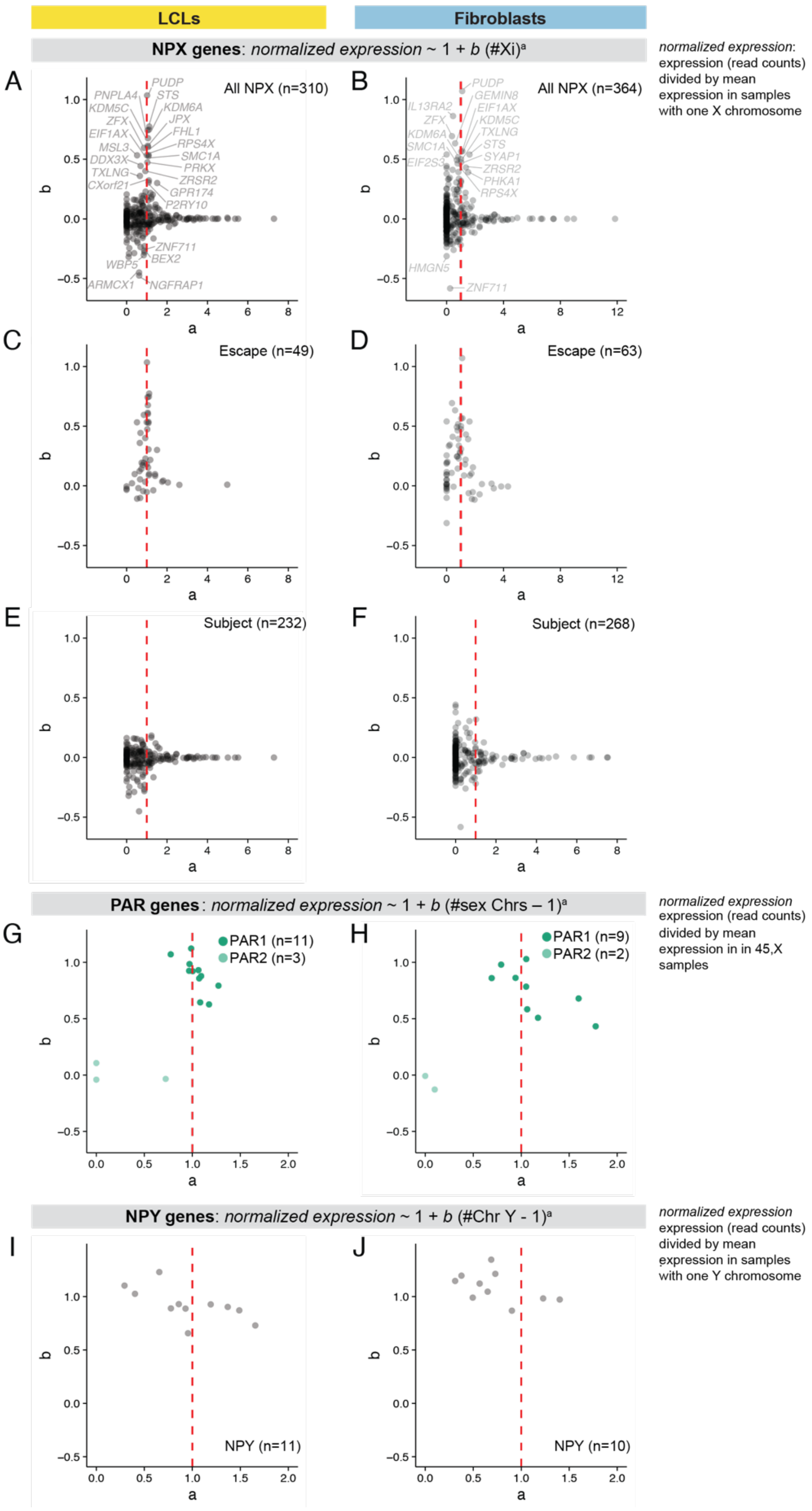

LCLs
Fibroblasts
NPX genes: normalized expression ~ 1 + b (#Xi)a
normalized expression: expression (read counts) divided by mean expression in samples with one X chromosome
A
All NPX (n=310)
B
All NPX (n=364)
C
Escape (n=49)
D
Escape (n=63)
E
Subject (n=232)
F
Subject (n=268)
PAR genes: normalized expression ~ 1 + b (#sex Chrs – 1)a
normalized expression expression (read counts) divided by mean expression in in 45,X samples
G
PAR1 (n=11)
PAR2 (n=3)
H
PAR1 (n=9)
PAR2 (n=2)
NPY genes: normalized expression ~ 1 + b (#Chr Y - 1)a
normalized expression expression (read counts) divided by mean expression in samples with one Y chromosome
I
NPY (n=11)
J
NPY (n=10)
a
b

**Figure S2. Linear functions are a good fit for modeling sex chromosome aneuploidy RNA-seq data.** For sex chromosome genes in LCLs and fibroblasts, power functions were fit (gray boxes) by non-linear least squares regression. Scatterplots show fitted values for the exponent parameter, *a*, and the coefficient, *b*, for each gene. *XIST* and genes for which the regression model did not converge are excluded from the plots. **(A-B)** Some NPX genes cluster near *a*=1, indicating their expression increases by a fixed amount for each additional X or Y chromosome. These correspond to genes previously annotated as escaping XCI **(C-D)**. Other NPX genes cluster near *a*=0 or *b*=0, indicating that they do not change in expression with additional copies of Xi, and correspond to genes previously annotated as subject to XCI (**E-F**). (**G-H**) PAR1 genes cluster near a=1, indicating their expression increases by a fixed amount for each additional sex chromosome, in contrast to PAR2 genes. (**I-J**) NPY genes also cluster near a=1, indicating their expression increases by a fixed amount for each Y chromosome.

**Figure S3.**

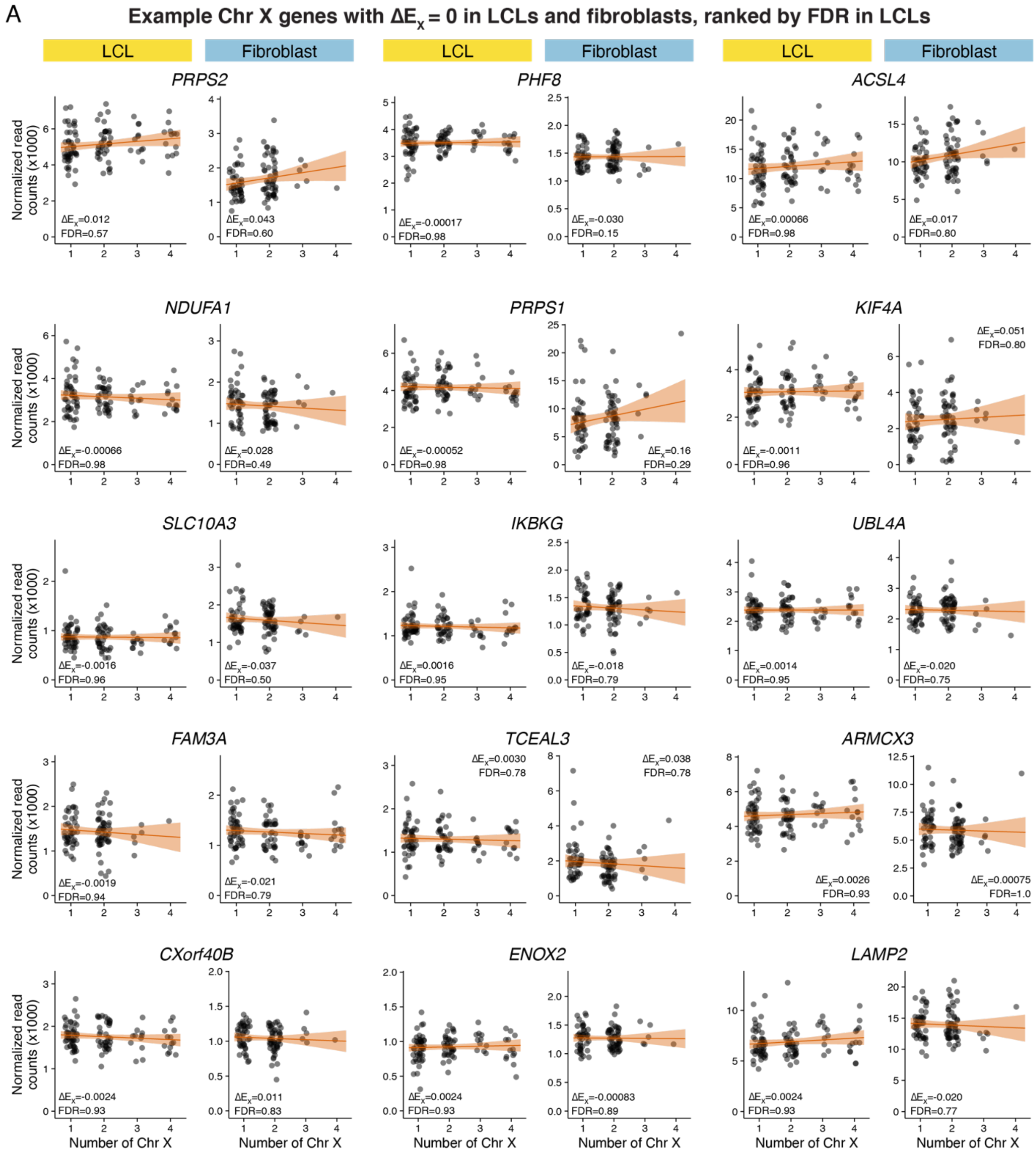
A
Example Chr X genes with ΔEx = 0 in LCLs and fibroblasts, ranked by FDR in LCLs
LCL
Fibroblast
LCL
Fibroblast
LCL
Fibroblast
PRPS2
ΔEx=0.012 FDR=0.57
ΔEx=0.043 FDR=0.60
PHF8
ΔEx=-0.00017 FDR=0.98
ΔEx=-0.030 FDR=0.15
ACSL4
ΔEx=0.00066 FDR=0.98
ΔEx=0.017 FDR=0.80
NDUFA1
ΔEx=-0.00066 FDR=0.98
ΔEx=0.028 FDR=0.49
PRPS1
ΔEx=-0.00052 FDR=0.98
ΔEx=0.16 FDR=0.29
KIF4A
ΔEx=-0.0011 FDR=0.96
ΔEx=0.051 FDR=0.80
SLC10A3
ΔEx=-0.0016 FDR=0.96
ΔEx=-0.037 FDR=0.50
IKBKG
ΔEx=0.0016 FDR=0.95
ΔEx=-0.018 FDR=0.79
UBL4A
ΔEx=0.0014 FDR=0.95
ΔEx=-0.020 FDR=0.75
FAM3A
ΔEx=-0.0019 FDR=0.94
ΔEx=-0.021 FDR=0.79
TCEAL3
ΔEx=0.0030 FDR=0.78
ΔEx=0.038 FDR=0.78
ARMCX3
ΔEx=0.0026 FDR=0.93
ΔEx=0.00075 FDR=1.0
CXorf40B
ΔEx=-0.0024 FDR=0.93
ΔEx=0.011 FDR=0.83
ENOX2
ΔEx=0.0024 FDR=0.93
ΔEx=-0.00083 FDR=0.89
LAMP2
ΔEx=0.0024 FDR=0.93
ΔEx=-0.020 FDR=0.77
Normalized read counts (x1000)
Number of Chr X

**Figure S3, continued.**

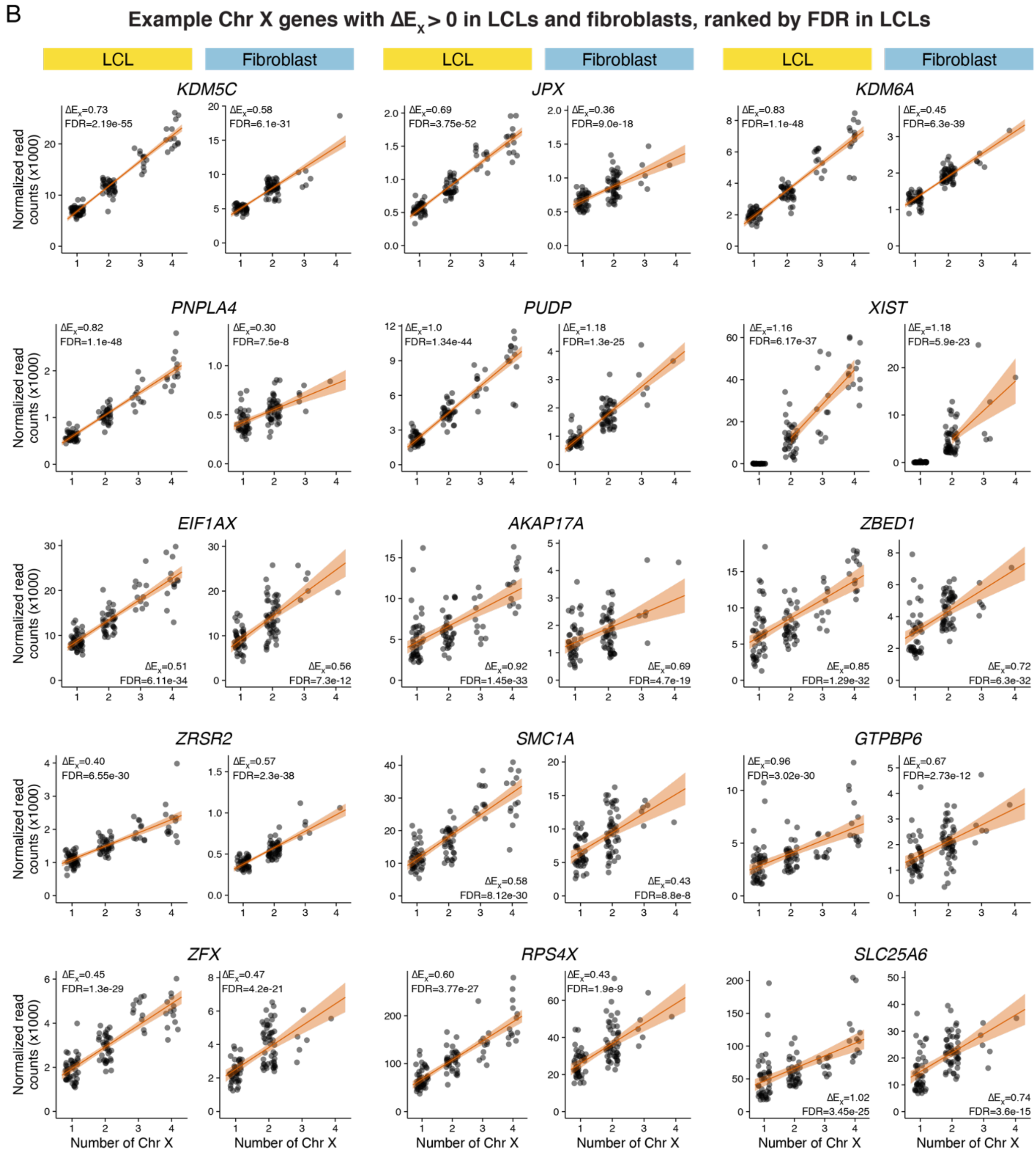

**Figure S3, continued.**

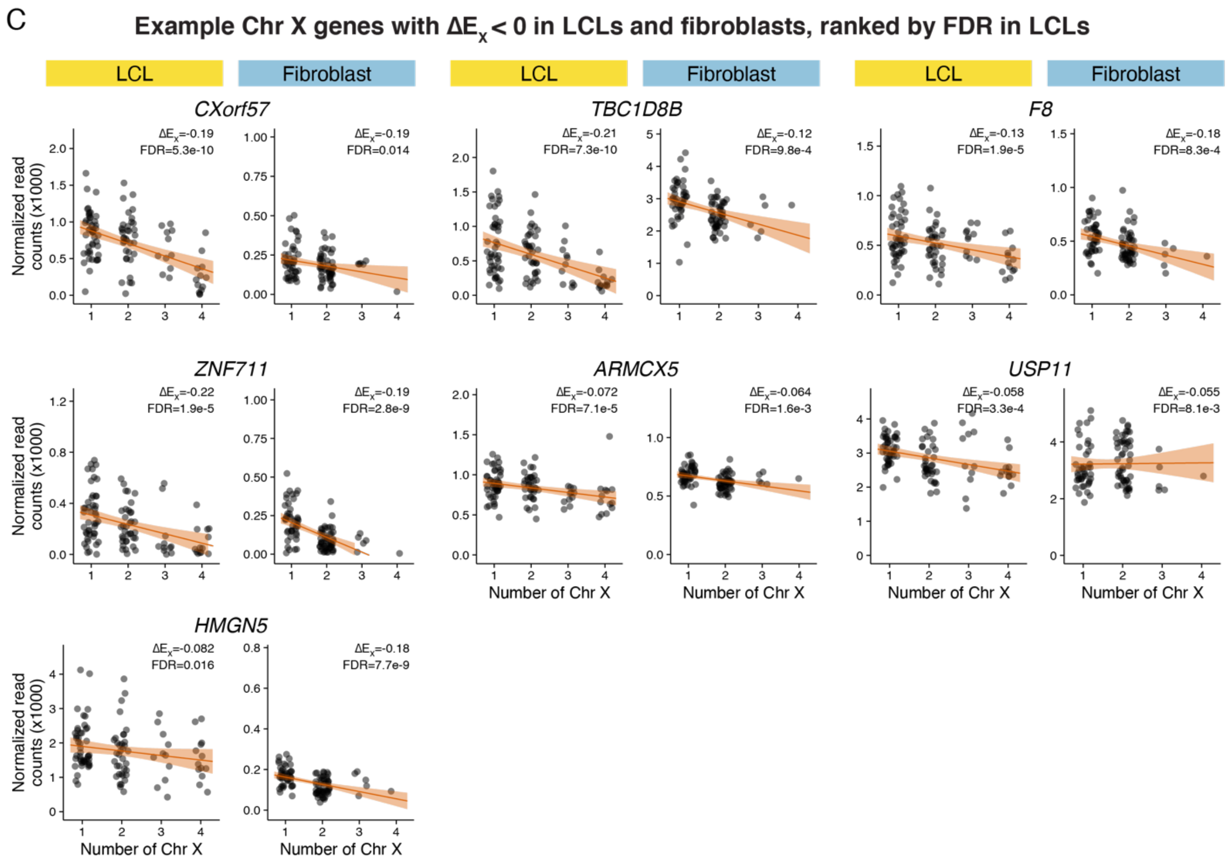

**Figure S3, continued.**

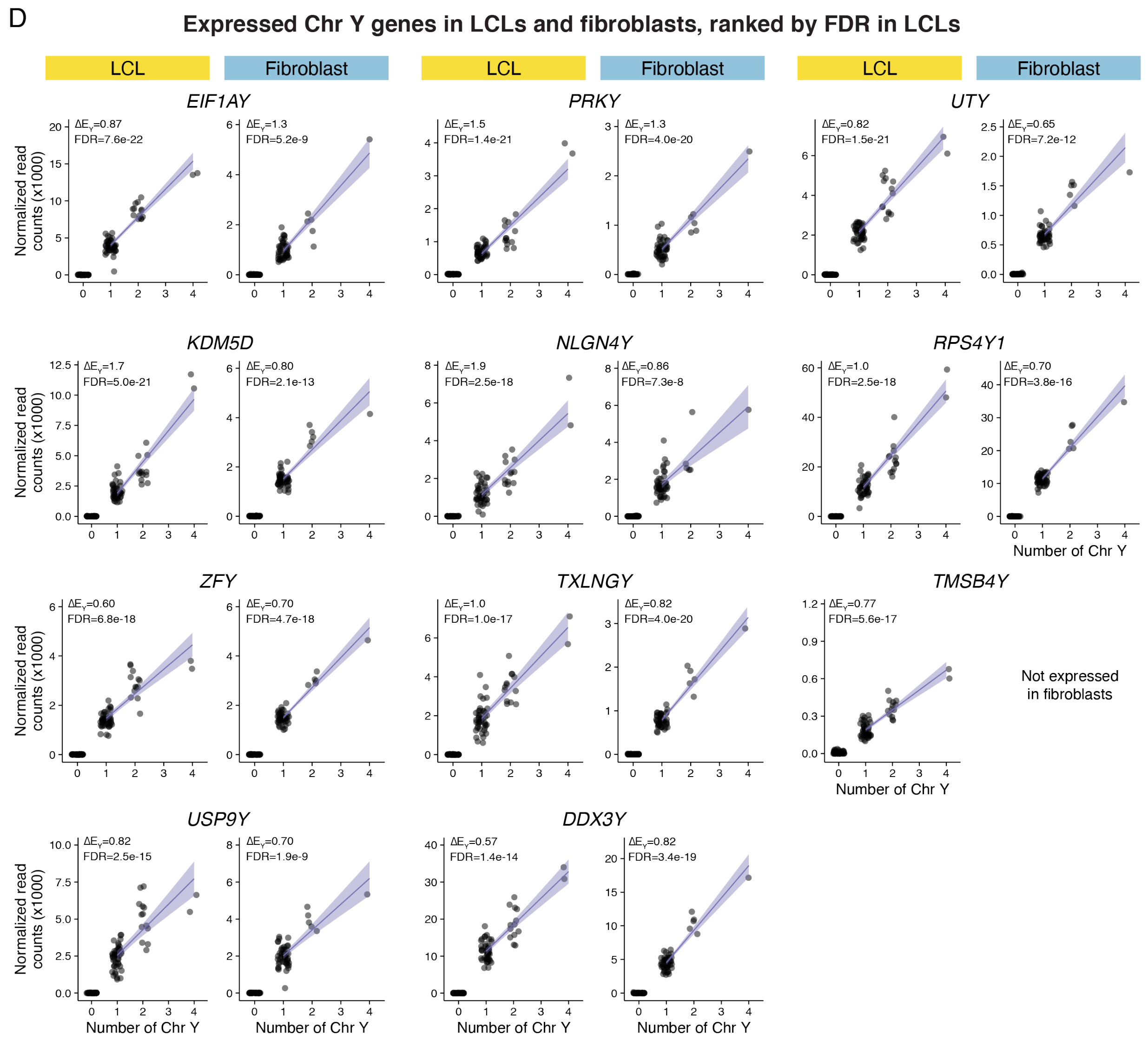

**Figure S3, continued.**

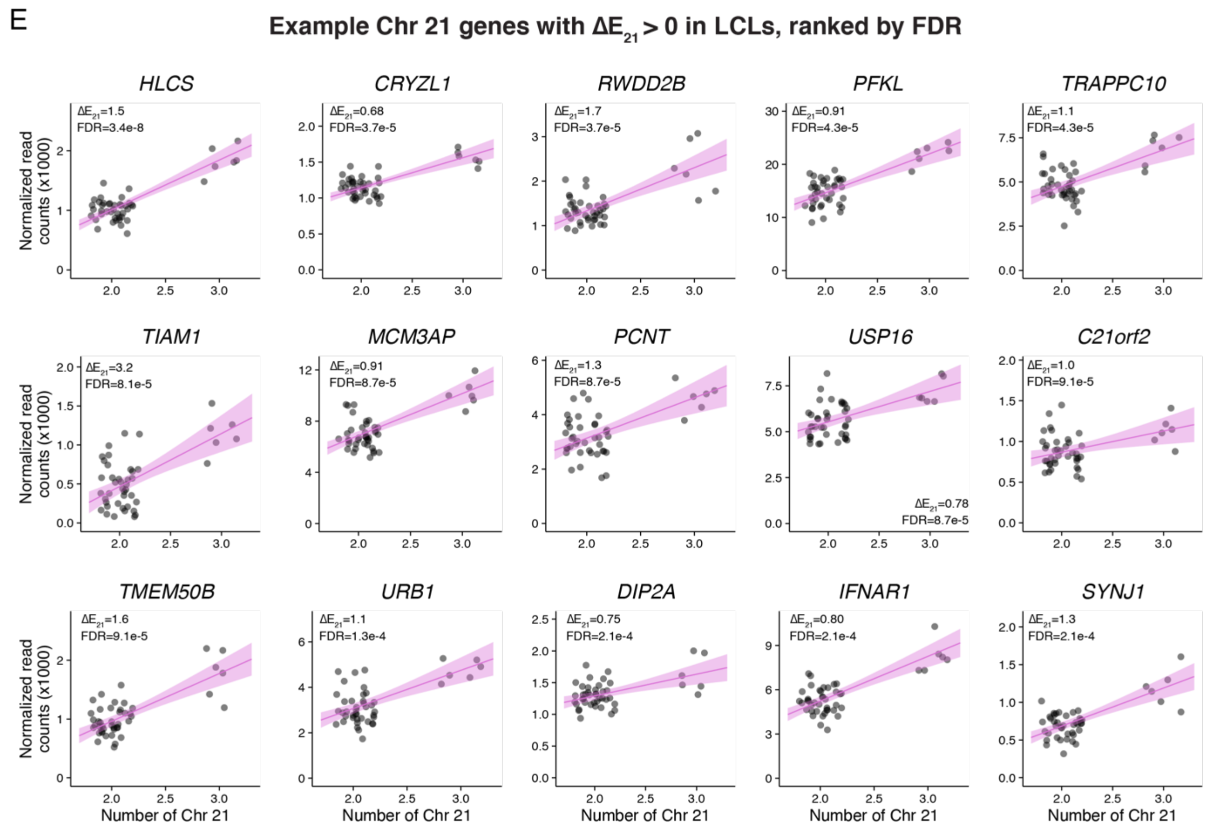


**Figure 3. Extended examples of ΔE values for Chr X, Y and 21 genes.**

Scatterplots and regression lines with confidence intervals for selected genes across Chr X, Y, or 21 copy number series. Adjusted p-values (FDR) <0.05 indicate that $\Delta E_X$ values are significantly different from zero. **(A)** Fifteen Chr X genes with $\Delta E_X$=0 in LCLs and fibroblasts. **(B)** Fifteen Chr X genes with $\Delta E_X$>0 in LCLs and fibroblasts. **(C)** Seven Chr X genes with $\Delta E_X$<0 in LCLs and fibroblasts. **(D)** All expressed Chr Y genes. **(E)** Fifteen Chr 21 genes with $\Delta E_{21}$>0.

**Figure S4.**

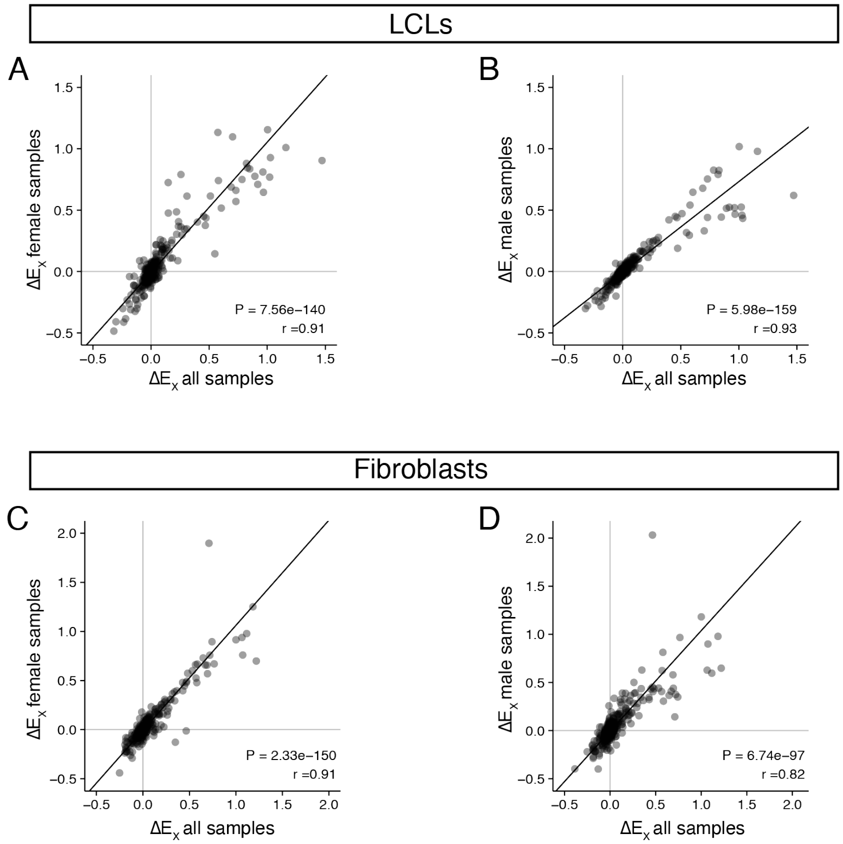


**Figure S4. $\Delta E_X$ values are similar when calculated with all samples, or with female (0 Y chromosomes) or male (1 Y chromosome) samples.** Scatterplots of all expressed genes in LCLs (A-B) or fibroblasts (C-D) showing $\Delta E_X$ values calculated using all samples (X-axes) or exclusively female samples (A, C Y-axes) or male samples (B, D Y-axes). Deming regression lines, Pearson correlations and P-values are indicated.

**Figure S5.**

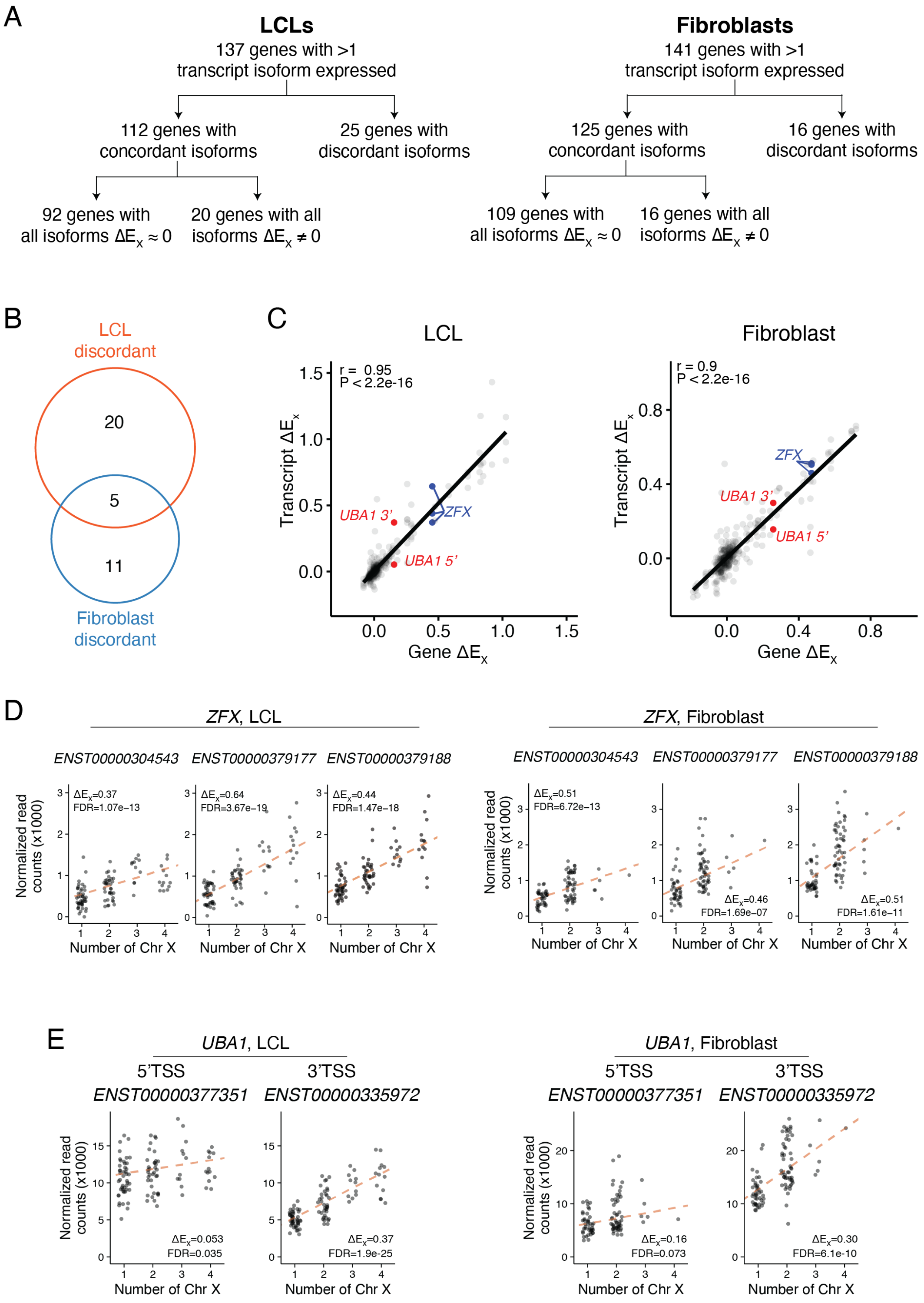
A
LCLs
137 genes with >1 transcript isoform expressed
112 genes with concordant isoforms
25 genes with discordant isoforms
92 genes with all isoforms ΔEx ≈ 0
20 genes with all isoforms ΔEx ≠ 0
Fibroblasts
141 genes with >1 transcript isoform expressed
125 genes with concordant isoforms
16 genes with discordant isoforms
109 genes with all isoforms ΔEx ≈ 0
16 genes with all isoforms ΔEx ≠ 0
B
LCL discordant
20
5
11
Fibroblast discordant
C
LCL
r = 0.95
P < 2.2e-16
Transcript ΔEx
Gene ΔEx
UBA1 3'
UBA1 5'
ZFX
Fibroblast
r = 0.9
P < 2.2e-16
D
ZFX, LCL
ZFX, Fibroblast
ENST00000304543
ENST00000379177
ENST00000379188
Normalized read counts (x1000)
Number of Chr X
ΔEx=0.37
FDR=1.07e-13
ΔEx=0.64
FDR=3.67e-19
ΔEx=0.44
FDR=1.47e-18
ΔEx=0.51
FDR=6.72e-13
ΔEx=0.46
FDR=1.69e-07
ΔEx=0.51
FDR=1.61e-11
E
UBA1, LCL
UBA1, Fibroblast
5'TSS
3'TSS
ENST00000377351
ENST00000335972
ΔEx=0.053
FDR=0.035
ΔEx=0.37
FDR=1.9e-25
ΔEx=0.16
FDR=0.073
ΔEx=0.30
FDR=6.1e-10

**Figure S5. Quantitative analysis of Xi contributions to transcript isoform expression (A)** Accounting of Chr X genes with multiple expressed transcript isoforms in LCLs and fibroblasts. Indicated here are the numbers of genes where i) all isoforms show $\Delta E_X \approx 0$, or ii) all isoforms show $\Delta E_X$ values significantly different from zero in the same direction (FDR<0.05), or iii) isoforms show discordant $\Delta E_X$ values. **(B)** Venn diagram shows the overlap of genes expressed in both LCL and fibroblasts that had isoforms with discordant $\Delta E_X$ values. **(C)** Scatterplots of each transcript isoform's $\Delta E_X$, compared with the gene's $\Delta E_X$ in LCLs and fibroblasts. Three *ZFX* isoforms are indicated with blue points; two *UBA1* isoforms indicated with red points. Pearson correlations (r) between isoforms and genes and P-values are indicated. **(D)** Scatterplots of expression of three *ZFX* isoforms across Chr X copy number in LCLs and fibroblasts showing that they consistently have $\Delta E_X > 0$. **(E)** Scatterplots of expression of two *UBA1* isoforms across Chr X copy number in LCLs and fibroblasts showing discordant $\Delta E_X$ values. Regression lines, $\Delta E_X$, and FDR are indicated.

**Figure S6.**

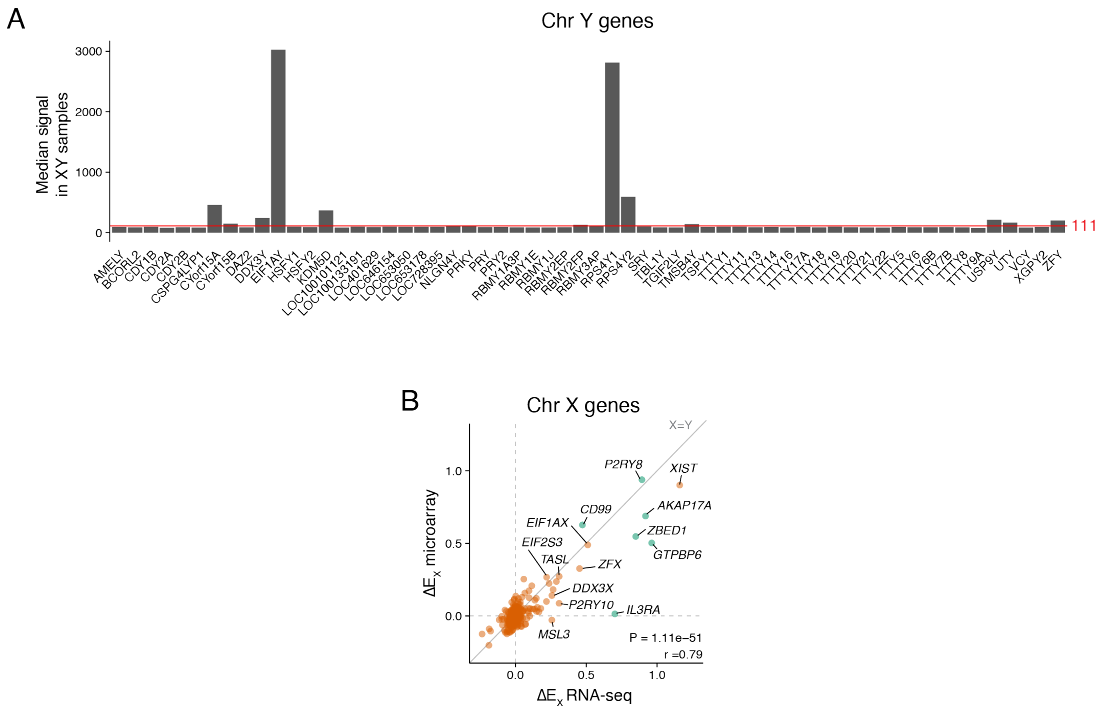


**Figure S6. Reanalysis of microarray data from LCLs of individuals with varying sex chromosome constitutions using linear modeling. (A)** Chr Y gene expression in microarray data from XY samples was assessed to set a minimum signal threshold of 111 (red line) for expressed genes, based on those expressed in LCL RNA-seq. **(B)** $\Delta E_X$ values in RNA-seq and microarray data for expressed Chr X genes were highly correlated. Deming regression line, Pearson correlation and P-value are indicated.

**Note S1**

***Divergent $\Delta E_X$ values for transcript isoforms of an X-linked gene***

To investigate the possible causes of discordant $\Delta E_X$ values across a gene's isoforms (25 genes in LCLs and 16 genes in fibroblasts), we asked whether any of the divergent isoforms had different transcription start sites (TSSs). two genes in LCLs (*ARHGEF6, BCOR*) and three genes in fibroblasts (*SH3KBP1, THOC2, UBA1*) had discordant isoforms with TSSs >500bp from each other, while the rest had very similar TSS locations. Of these genes, *UBA1* is the only gene where its flanking genes had different $\Delta E_X$ values. Moreover, the isoform $\Delta E_X$ values correlate with those of the flanking genes: the *UBA1* transcript from the 5' start (ENST00000377351) and upstream gene *RBM10* had $\Delta E_X$ values near zero, while the *UBA1* transcript from the 3' start (ENST0000033972) and downstream gene *CDK16* had significantly positive $\Delta E_X$ values. Analysis of ENCODE data reveals a strong CTCF binding site between the two *UBA1* TSSs in LCLs and fibroblasts, and evidence of divergent three-dimensional interactions, suggesting that the two start sites reside in distinct topologically associated domains along with their neighboring genes.

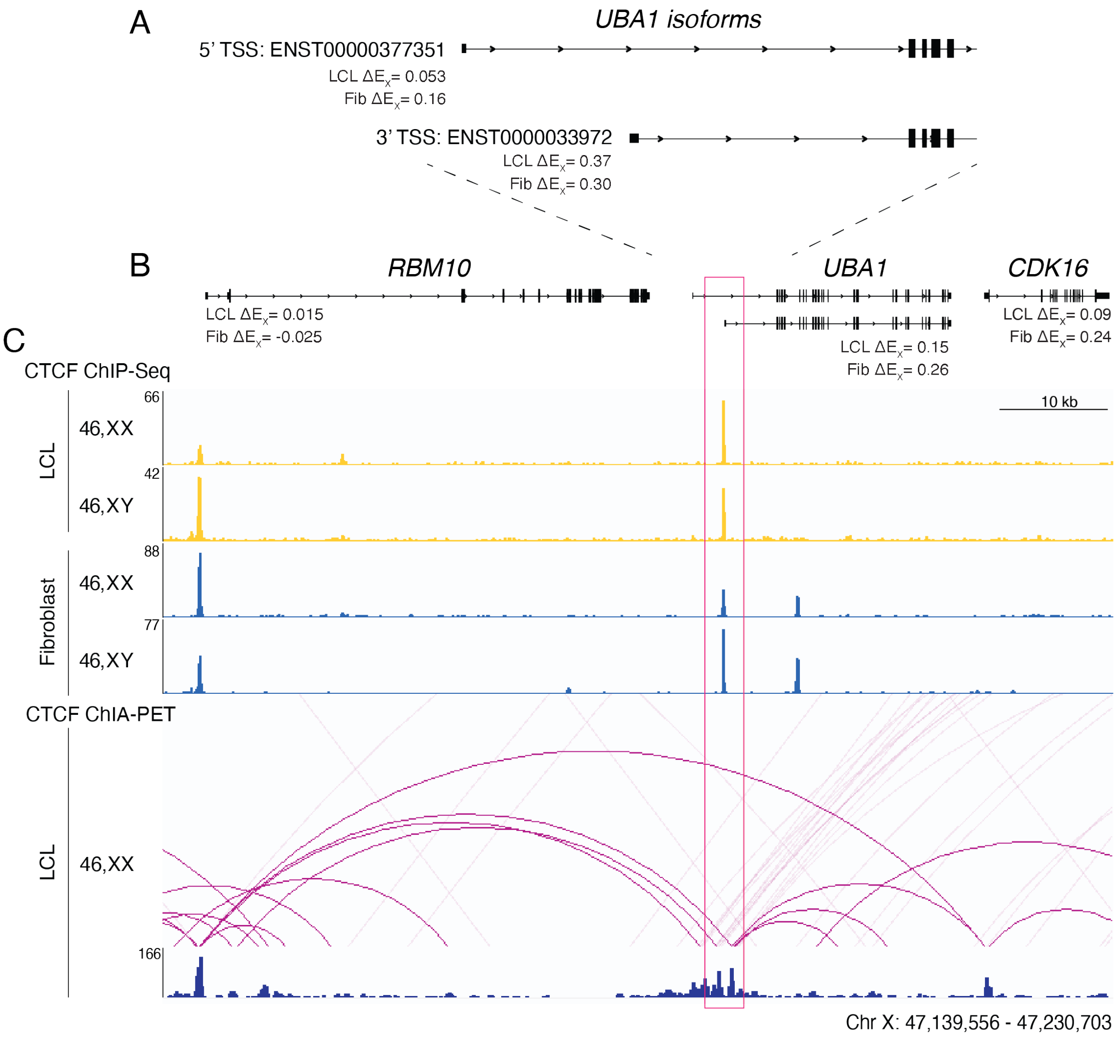


**ChIP-seq and ChIA-PET for CTCF reveal a domain boundary between two *UBA1* isoform transcription start sites.** **(A)** Transcript models for *UBA1* transcript isoforms. Isoform-level $\Delta E_X$ values are listed. **(B)** Gene models for *UBA1* and flanking genes *RBM10* and *CDK16.* Gene-level $\Delta E_X$ values are listed. **(C)** CTCF ChIP-seq signal tracks (fold-change over background) for two LCL and two fibroblast cell lines show a strong CTCF peak between the two *UBA1* isoform transcription start sites. Interaction and signal tracks (fold-change over background) are shown for CTCF ChIA-PET. Red box indicates boundary region.

Three previous studies corroborate these results: 1) in human-mouse hybrid cell lines, the 5’ TSS was only active in lines containing Xa, while the 3’ TSS was active in lines containing Xa or Xi (Goto and Kimura, 2009); 2) 3’ TSS transcripts were more abundant in 46,XX than 46,XY cells, while 5’ TSS transcripts showed no such difference (Chen et al., 2016) and 3) DNA methylation studies revealed hypermethylation of the 5’ but not the 3’ TSS in XX cells from human, chimp, and horse (Balaton et al., 2021). Together, this publicly available data and our own analysis shows that *UBA1* is a unique example of a gene with transcription start sites that are divergently expressed from Xi.

**Note S2**

**Identifying cell lines with skewed X chromosome inactivation and calculating allelic ratios from RNA-seq data**

46,XX individuals typically have a mixture of cells in which either the maternal or paternal X chromosome is active. Because of this, Chr X gene expression measured in bulk RNA-seq data from a mixed population of cells typically appears biallelic. However, in cells with skewed XCI – where most cells in the culture have the same Xa – expression from Xa only or both Xa and Xi can be distinguished.

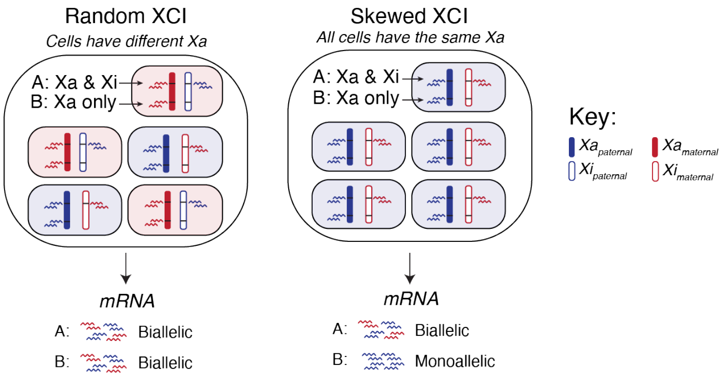


**Identifying skewed XCI in RNA-seq data.** Schematic showing allele-specific expression results in 46,XX cell cultures with random or skewed XCI. In mRNA extracted from the random XCI culture, all genes will appear biallelic, while in mRNA extracted from the skewed XCI culture, genes expressed only from Xa appear monoallelic, while genes expressed from both Xa and Xi appear biallelic.

The overall workflow is summarized here, with each step described in detail below.

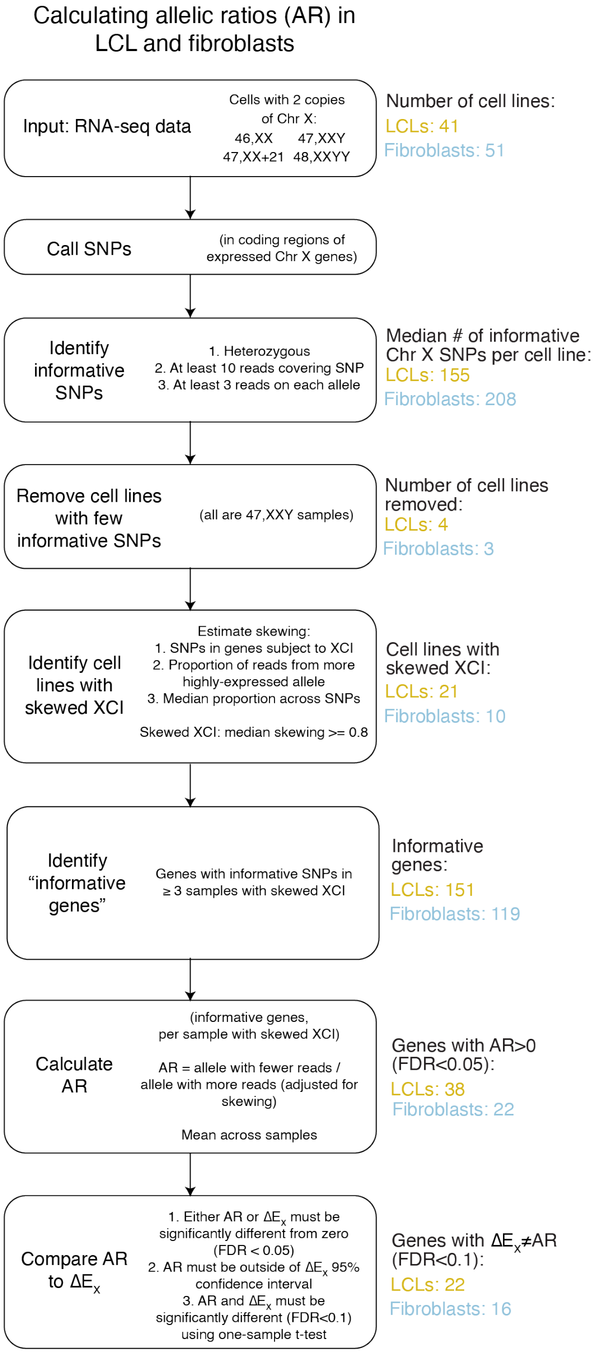


**Analytical workflow.** Each box indicates a step in the analytical pipeline described in more detail below. To the right of each box are summaries of results at each step.

We first called SNPs in samples with two copies of Chr X (46,XX, 46,XX,+21, 47,XXY, 48,XXYY; **Methods**) to identify those that were heterozygous, and could therefore distinguish between two alleles (SNP data provided in **Tables S6C-D**). Of the 41 LCL and 51 fibroblast samples with two copies of Chr X, we removed 8 (4 LCLs and 4 fibroblasts) because they had few heterozygous SNPs identified on Chr X. These samples all had the karyotype 47,XXY and had a similar number of heterozygous SNPs on Chr 8 compared to other samples, indicating that there is not a problem with these sample, but rather that they likely have two copies of the same Chr X inherited through a maternal meiotic non-disjunction.

For samples with sufficient heterozygous Chr X SNPS, we proceeded to identify samples with skewed XCI. We first curated a list of genes that were previously shown to only be expressed from Xa ("silenced" on Xi) and should appear monoallelic in cell lines with skewed XCI. The "Xa only" list was compiled from genes classified as "silenced" in recently-published studies (see **Methods** and **Table S6A**) and was restricted to those with non-significant changes with Chr X dosage in our dataset ($\Delta E_X$<0.05 and FDR>0.5). The lists of genes used in this analysis are in **Table S6E.** For SNPs in "Xa-only" genes, we calculated the "skewing coefficient" by dividing the allele with the higher read count by the total number of reads for that SNP, that is, the fraction of reads accounted for by one allele. This value can range from 0.5 (equal reads from two alleles) to 1 (all reads are from one allele). For each sample, we calculated the median skewing coefficient across all SNPs in "Xa-only" genes and samples with at least 0.8 median skewing coefficients were considered skewed – 11 LCL samples and nine fibroblast samples.

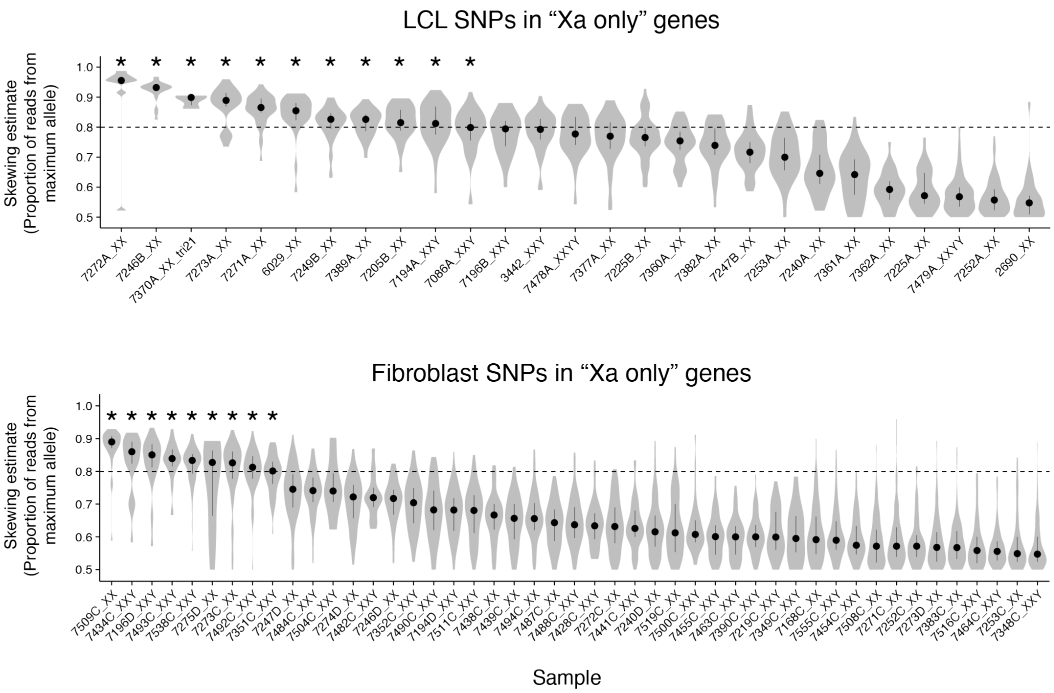


**Skewing analysis for SNPs in Chr X genes previously identified as expressed only from Xa.** Samples with >5 heterozygous SNPs in “Xa only” genes are ranked by the median skewing estimate across SNPs in “Xa only” genes. Asterisks, samples we considered as having skewed XCI (skewing estimate ≥ 0.8).

Some samples (10 LCL samples and one fibroblast culture) had ≤5 SNPs in “Xa only” genes, but many SNPs in other Chr X genes. Since we required that a SNP be heterozygous to be considered in our analyses, any cell line with 100% skewed XCI would display an absence of heterozygous SNPs identified in Xa only genes, since they should be monoallelically expressed. In these samples we identified heterozygous SNPs in Chr X genes that were not “Xa only”, meaning that there are not two identical copies of Chr X, but rather that that these samples were highly-skewed.

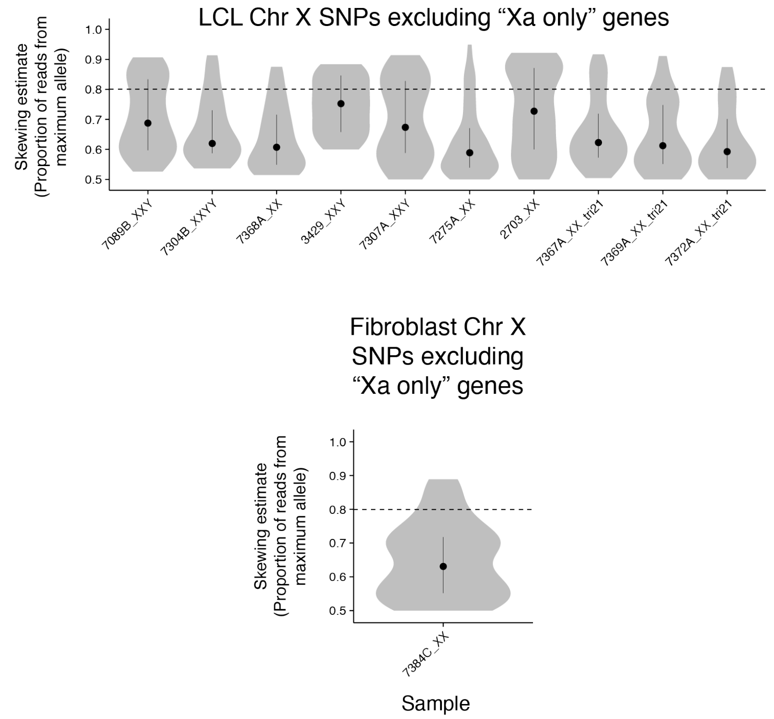


**SNPs in Chr X genes (excluding “Xa only genes”) for highly-skewed samples.** Samples with ≤5 SNPs in “Xa only” genes have heterozygous SNPs in other Chr X genes.

Between the samples with skewed and highly-skewed XCI, we identified 21 LCL and 10 fibroblast samples. To ensure that our skewing estimates for Chr X are not reflective of a systematic issue within individual samples, we analyzed SNPs on Chr 8, which has a comparable number of expressed genes. We invariably observed similar numbers of reads from the two alleles of a single locus.

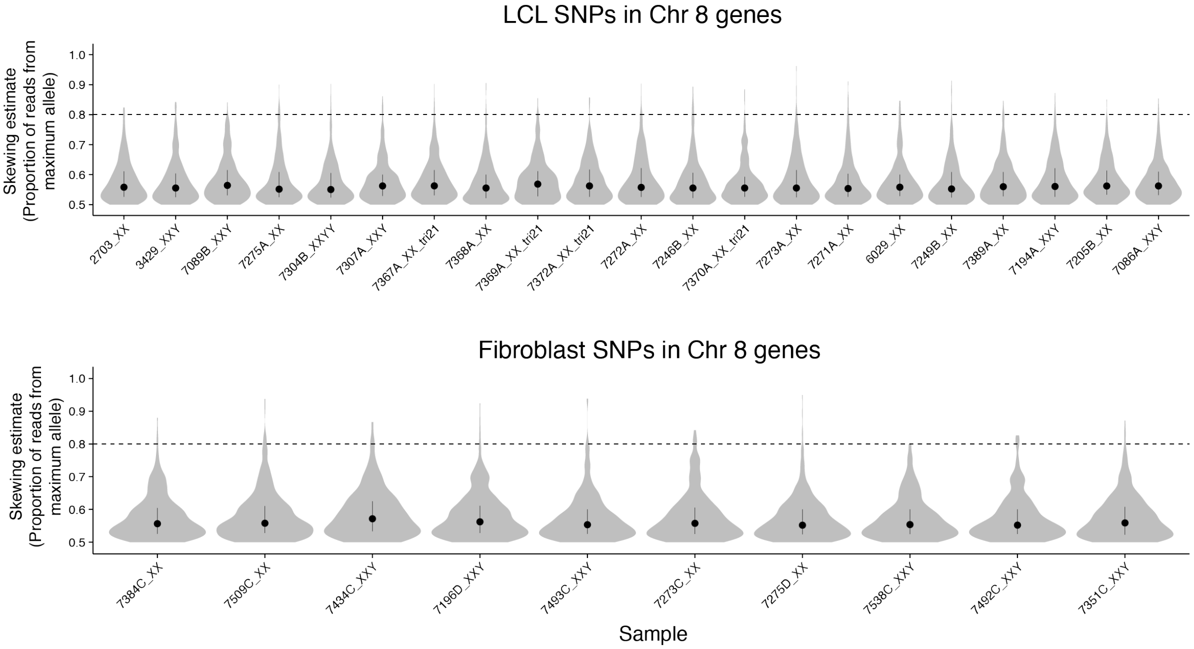


**Skewing analysis for SNPs in Chr 8 genes.** Skewing analysis was performed using SNPs in expressed genes on Chr 8 on samples with skewed XCI. Note uniform skewing estimates of <0.6 across LCL and fibroblast samples, indicating relatively equal expression of each allele.

Following identification of samples with skewed XCI, we identified "informative genes," requiring informative SNPs in at least 3 samples with skewed XCI, resulting in 151 genes in LCLs and 119 in fibroblasts. We then calculated the allelic ratio (AR) at each SNP by dividing the allele with fewer reads (assumed to represent Xi) by the allele with more reads (assumed to represent Xa), adjusting for the level of skewing in each sample using the median skewing estimate (skewing coefficient; **Methods; Table S6B**).

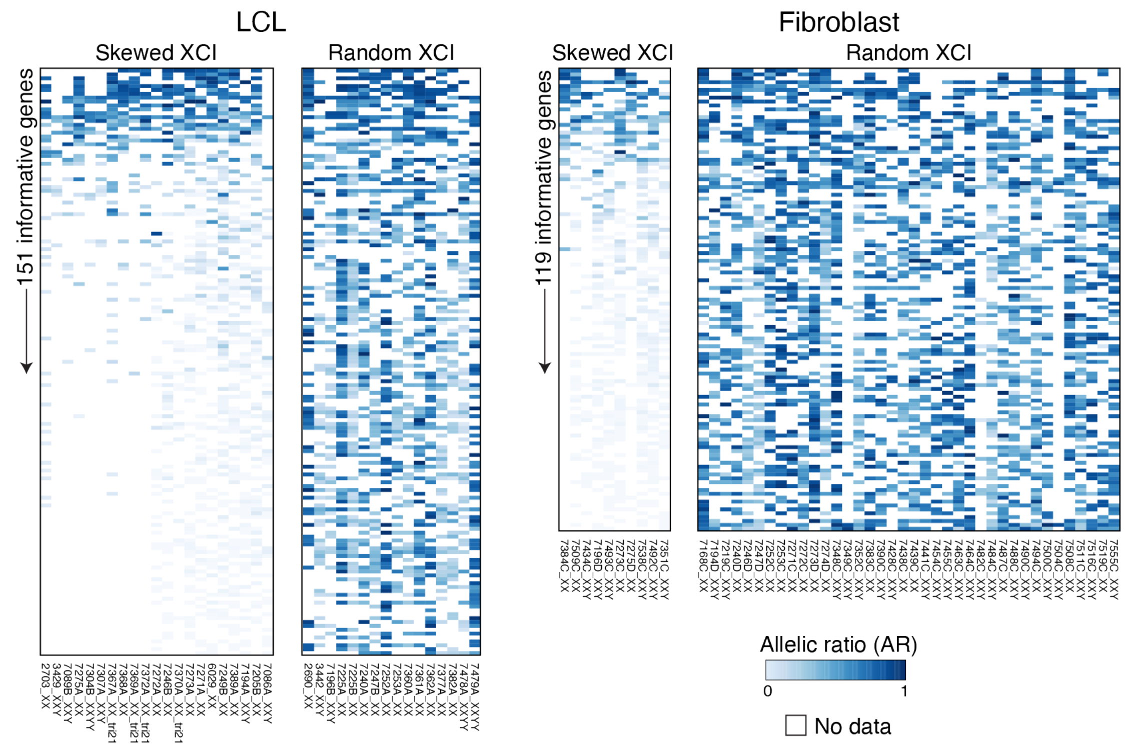


**Allelic ratio of informative genes in samples with skewed or random XCI.** Heatmaps showing adjusted allelic ratio data for genes with data in at least 3 samples with skewed XCI in LCLs or fibroblasts. Genes are ordered by the average adjusted AR across samples with skewed XCI. Raw AR values for samples that did not reach our 0.8 median skewing thresholds – termed random XCI – are shown for comparison. Note that genes with high and low allelic ratios can be distinguished in samples with skewed XCI, but not in samples with random XCI.

We validated these AR values by comparing them to published AR values computed in independent studies and found high correlations in each case (Cotton et al., 2013, Tukiainen et al., 2017, Garieri et al., 2018, Sauteraud et al., 2021).

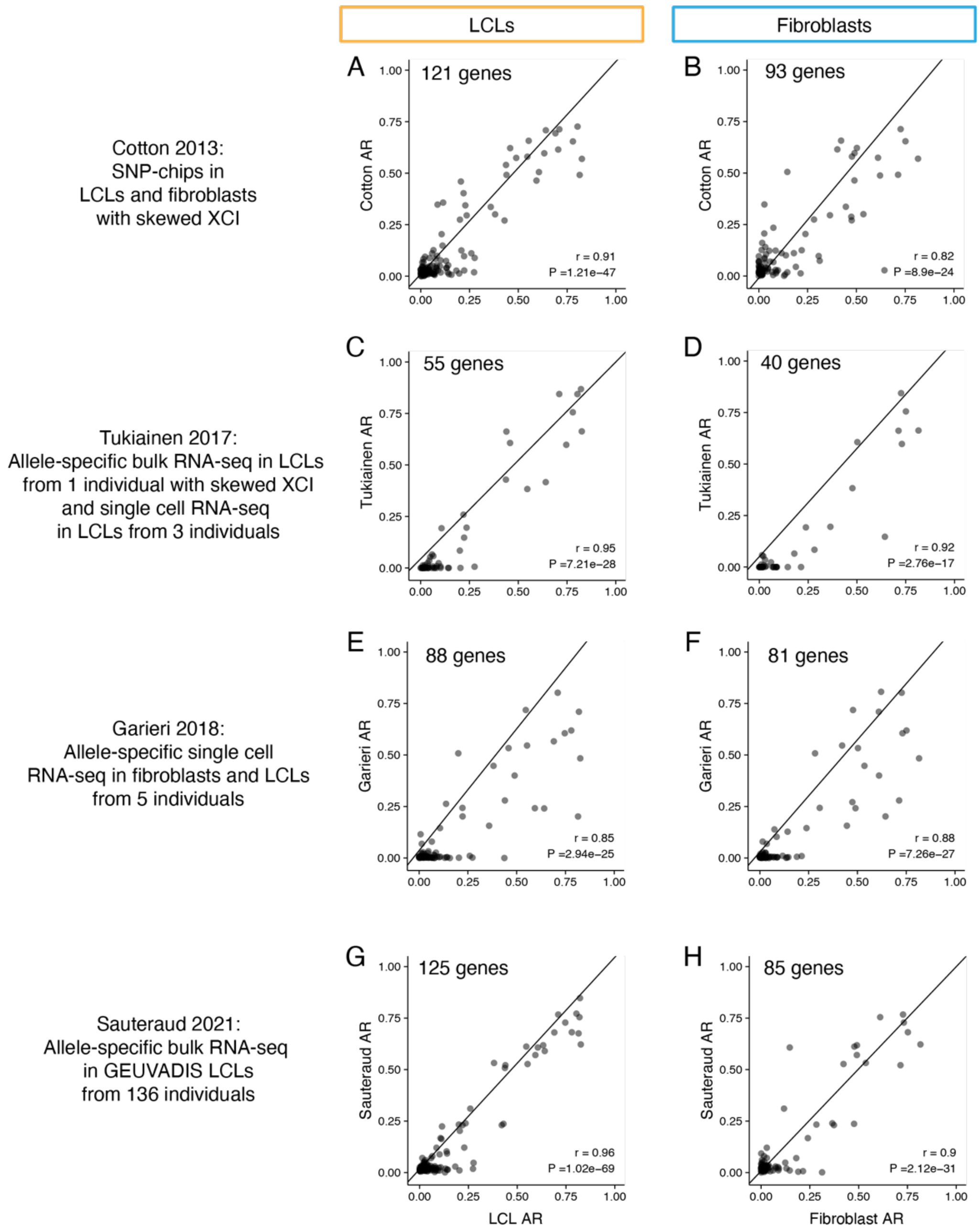


**Comparison of allelic ratios with published datasets.** Scatterplots of the AR calculated in this study versus published studies for informative genes included in both studies. Deming regression lines, Pearson correlation and P-values are indicated.

It is possible that by including the “highly skewed” samples, that the allelic ratios could be over-estimated. To test this, we repeated the analysis excluding highly-skewed samples, since the skewing coefficient could be stringently determined. This removed some informative genes from the analysis, but did not significantly change the AR levels (**Table S6F**).

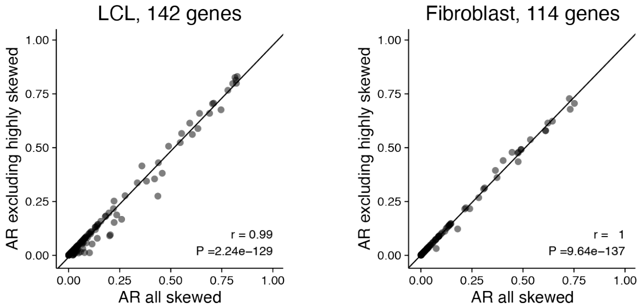


**Comparison of AR values calculated using all skewed samples or without the highly-skewed samples.** Scatterplots of AR calculated using the two sample sets. Deming regression lines, Pearson correlation and P-values are indicated.

**Table S1.** Metadata for euploid and aneuploid samples with RNA-sequencing.

**Table S2.** Linear regression results for expressed PAR and NPX genes

**Table S3.** Linear regression results for expressed PAR and NPX transcript isoforms

**Table S4.** Linear regression results for expressed NPY genes

**Table S5.** Linear regression results for expressed Chr 21 genes

**Table S6.** Comparison of $\Delta E_X$ values to XCI status calls from published studies and allelic ratios from this study

**Table S7.** Expression constraint metrics for PAR1 and NPX genes

*All supplemental tables are provided as .xlsx spreadsheets.*
